# An epigenetic breeding system in soybean for increased yield and stability

**DOI:** 10.1101/232819

**Authors:** Sunil Kumar Kenchanmane Raju, Mon-Ray Shao, Robersy Sanchez, Ying-Zhi Xu, Ajay Sandhu, George Graef, Sally Mackenzie

## Abstract

Epigenetic variation has been associated with a wide range of adaptive phenotypes in plants, but there exist few direct means for exploiting this variation. RNAi suppression of the plant-specific gene, *MutS HOMOLOG1* (*MSH1*), in multiple plant species produces a range of developmental changes accompanied by modulation of defense, phytohormone, and abiotic stress response pathways. This *msh1-conditioned* developmental reprogramming is retained independent of transgene segregation, giving rise to transgene-null ‘memory’ effects. An isogenic memory line crossed to wild type produces progeny families displaying increased variation in adaptive traits that respond to selection. This study investigates amenability of the *MSH1* system for inducing epigenetic variation in soybean that may be of value agronomically. We developed epi-line populations by crossing with *msh1*-acquired soybean memory lines. Derived soybean epi-lines showed increase in variance for multiple yield-related traits including pods per plant, seed weight, and maturity time in both greenhouse and field trials. Selected epi-F_2:4_ and epi-F_2:5_ lines showed an increase in seed yield over wild type. By epi-F_2:6_, we observed a return of MSH1-derived enhanced growth back to wild type levels. Epi-populations also showed evidence of reduced epitype-by-environment (e × E) interaction, indicating higher yield stability. Transcript profiling of the soybean epi-lines identified putative signatures of enhanced growth behavior across generations. Genes related to cell cycle, abscisic acid biosynthesis, and auxin-response, particularly SMALL AUXIN UP RNAs (SAURs), were differentially expressed in epi-F_2:4_ lines that showed increased yield when compared to epi-F_2:6_. These data support the potential of *msh1*-derived epigenetic variation in plant breeding for enhanced yield and yield stability.

## INTRODUCTION

Plants respond to changing environments through phenotypic plasticity that derives from both genetic and epigenetic factors (Bossdorf et al., 2010; Kooke et al., 2015). Epigenetic variation can, to some extent, be monitored via cytosine DNA methylation repatterning (Becker et al., 2011; Schmitz et al., 2011) that can be transgenerationally heritable (Quadrana and Colot, 2016). Arabidopsis epigenetic recombinant inbred lines (epiRILs), derived from crossing wild type *Col-0* with *met1* or *ddm1* DNA methylation mutants, show segregation and heritability of novel methylation patterns together with phenotypic diversity (Johannes et al., 2009; Reinders et al., 2009; Roux et al., 2011). The epiRILs show variation in biomass productivity, especially when challenged with weed competitors and biotic stress, driven partly by complementarity among epigenotypes (Latzel et al., 2013). Variation in complex traits like flowering time and root length is also influenced by epigenetic variation of segregating DNA methylation changes (Cortijo et al., 2014). These observations advance the hypothesis that induced epigenetic variation can be exploited effectively for selection in crop improvement.

*MutS HOMOLOG1* (*MSH1*) is a plant-specific homolog of the bacterial DNA repair gene *MutS* (Abdelnoor et al., 2003). MSH1 is a nuclear-encoded protein that is dual-targeted to mitochondria and plastids, and depletion of MSH1 influences both mitochondrial and plastid properties (Xu et al., 2011). In Arabidopsis *msh1* T-DNA insertion lines, phenotypes include leaf variegation, reduced growth rate, delayed flowering, extended juvenility, altered floral morphology, aerial rosettes, and enhanced secondary growth (Xu et al., 2012). These mutants also show tolerance to heat, high light and drought stress (Shedge et al., 2010; Virdi et al., 2016; Xu et al., 2011). These pleiotropic phenotypes are largely attributed to depletion of MSH1 from plastids, evidenced by hemi-complementation analysis (Xu et al., 2012), and the *msh1*-triggered plastid changes condition genome-wide methylome repatterning (Virdi et al., 2015). Similarly, detailed transcriptome analysis of *msh1* mutants reveals wide-ranging changes in gene expression related to defense response, abiotic stress, MAPK cascade, circadian rhythm, and phytohormone pathways (Shao et al., 2017).

RNAi suppression of *MSH1* in monocot and dicot species produces an identical range of developmental phenotypes (de la Rosa Santamaria et al., 2014; Xu et al., 2012; Yang et al., 2015). The altered phenotypes are somewhat attenuated but stable after segregation of the RNAi transgene, producing *msh1* ‘memory’. In sorghum, crossing *msh1* memory lines with isogenic wild type gives rise to enhanced vigor phenotypes that appear to respond to selection in small-scale studies (de la Rosa Santamaria et al., 2014). In tomato, *msh1*-derived vigor phenotypes are heritable in greenhouse and field conditions, graft transmissible and obviated by treatment with 5-azacytidine, further implicating DNA methylation in this phenomenon (Yang et al., 2015).

Soybean (*Glycine max* (L.)*Merr.*) is the most widely grown legume in the world, second only to grasses in economic importance. Synergistic interactions between advances in breeding and agronomic practices have steadily increased soybean yields in the past century (Rowntree et al., 2013). Further improvement will face challenges from climate instability and limited genetic diversity, which calls for the implementation of novel tools and methodologies to benefit soybean performance over a broad range of environments (Rincker et al., 2014). In this study, we used the well-known soybean variety ‘Thorne’ (McBlain et al., 1993) to investigate amenability of the *MSH1* system in exploiting epigenetic breeding potential. Greenhouse and large-scale multi-location field trials showed enhanced yield in selected F_2:4_ and F_2:5_ epi-lines. We document tapering of *msh1*-derived vigor in these lines by F_2:6_, and show evidence of buffering effects in epi-populations across environments, thus reducing epitype-by-environment interaction and possibly stabilizing yield across locations. Transcriptome studies of epi-lines from F_2:4_, F_2:5_ and F_2:6_ generations revealed genes and pathways that participate in the *msh1*-derived enhanced growth and its waning by later generations.

## RESULTS

### *MSH1* suppression in soybean induces a characteristic pleiotropic phenotype that persists after transgene segregation

RNAi suppression of *MSH1* in soybean produces phenotypic changes that include reduced growth rate, male sterility, enhanced branching, and altered leaf and floral morphology (Fig 1A), similar to earlier reports in Arabidopsis, tomato, and tobacco (Sandhu et al., 2007; Xu et al., 2011). Severely affected plants grow slower than wild type (Fig 1B), and show delayed flowering, extended juvenility and enhanced branching. The soybean *msh1*-RNAi T_0_ population did not produce visible variegation and/or male sterility, although 10 – 20% of progeny from these lines (T_1_) showed wrinkled and puckered leaves. Almost 50% of the T_1_ plants were semisterile, with increased flower drop and partially filled or empty seed pods. In subsequent generations, plants displayed a variable range of phenotypic severity.

**Figure 1.**
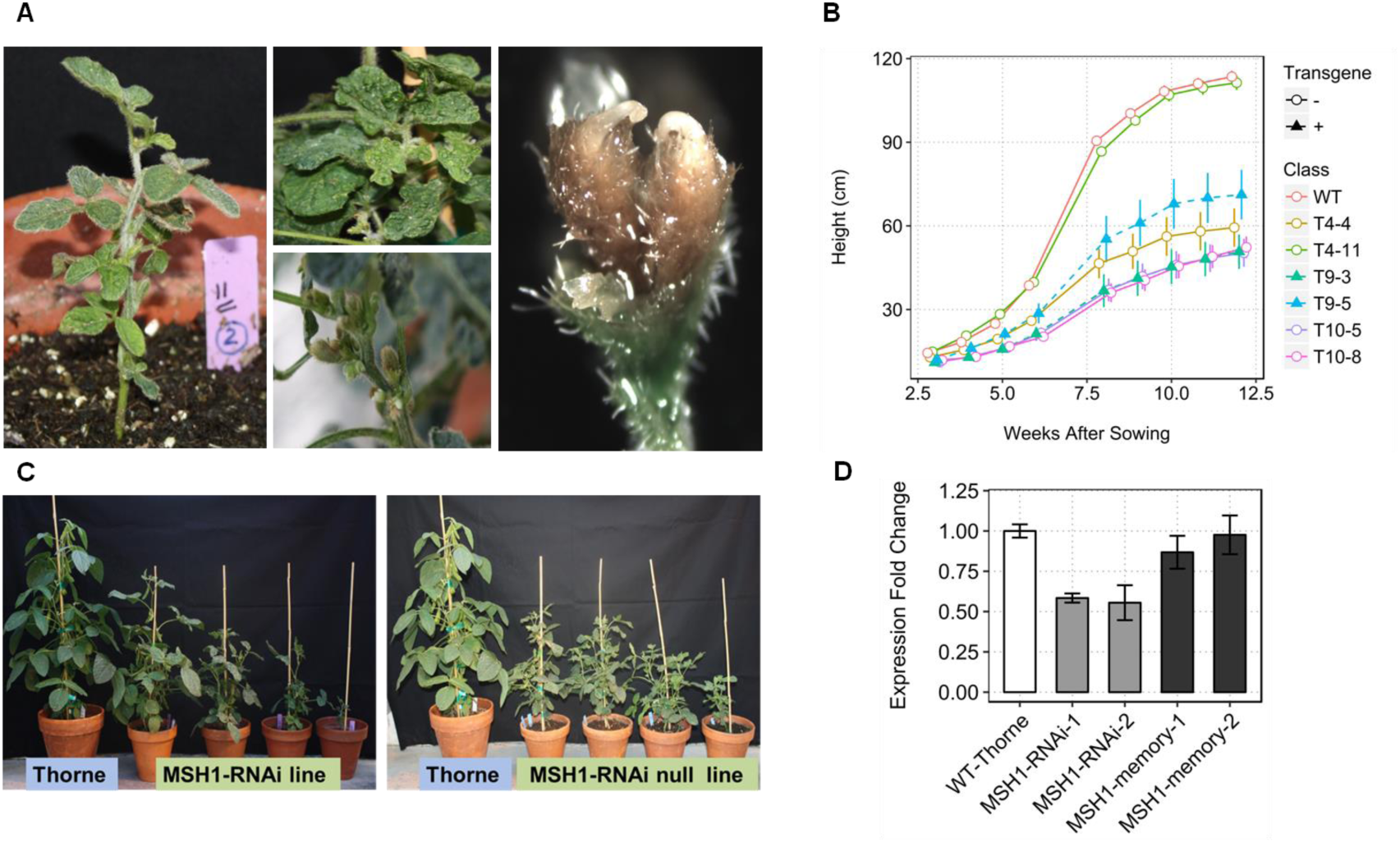
Characteristic phenotypes of *MSH1* suppression in soybean. **A)** Characteristic *MSH1*-RNAi phenotypes, dwarfing, wrinkled leaves, alterations of pod emergence and altered floral morphology showing flower with two stigmas. **B)** Growth-curve based on plant height in cm (measured weekly after 3 weeks of sowing) showing *MSH1*-RNAi and transgene-null *msh1* memory lines with reduced growth rate and higher variability within lines compared to wild type. **C)** Similar range in plant height and leaf morphology variation from T7 *MSH1*-RNAi (with transgene) and T10 *msh1* memory (without transgene) compared to wild ^t^yp^e^. **D)** Gene expression profiling of *MSH1*-RNAi and *msh1* memory lines for suppression of *MSH1* transcript level. Gene expression normalized to soybean actin levels and error bars represent SEM from three biological replicates.

Following transgene segregation, a proportion of progeny retained their acquired phenotypes of dwarfing, delayed flowering and altered leaf morphology for seven self-pollinated generations tested to date (Fig 1C). The transgene-null lines, retaining altered phenotype while restored in *MSH1* transcript levels (Fig 1D), comprise the memory lines used in this study. Memory lines were classified based on their phenotype into intermediate (i) and extreme (e) designated *i*MSH1** and *e*MSH1** respectively, while the remaining did not show any visible *MSH1* phenotype and were categorized n*MSH1* (Fig S1A).

### Transcript profiling of soybean *msh1*-RNAi lines shows correspondence of gene expression changes with phenotype severity

To evaluate the association of transcriptome changes with severity of *MSH1* phenotype, two soybean MSH1-RNAi lines (transgene positive) differing in their phenotype severity were assayed by gene expression profiling with the Affymetrix Soybean Genome Array (GPL4592) (Xu et al., 2011). We used a stringent cutoff (p-value < 0.05 and |log2(value)| > 1) to call differentially expressed genes (DEGs) relative to wild type controls.

The severe phenotype plants showed differential expression of 2589 genes, whereas mild phenotype plants showed 154 DEGs, 114 of which were shared in common (Fig S2A). Both classes had far more up-regulated genes, with severe showing 1656 up-regulated and 933 down-regulated genes, and mild showing 145 up-regulated and only nine down-regulated genes (Fig S2B, Table S1). The photosystem II-related genes PsbP-like 2, PsbQ-like 2 and PS II reaction center PSB28 (Glyma.10g290700, Glyma.12g215100, Glyma.13g127200) were significantly down-regulated, similar to what was described in Arabidopsis *msh1* mutants (Shao et al., 2017). We also observed significant down-regulation of histone H3, H4 and H2B.3 proteins (Glyma.15g032300, Glyma.20g083800, and Glyma.12g179100), consistent with plant stress response (Logemann et al., 1995).

Gene Ontology (GO) analysis with AgriGO (Du et al., 2010) classified differences between the two phenotypic classes. While mild-phenotype plants showed predominantly abiotic stress response, severe-phenotype plants were more broadly affected in phytohormone, defense, immune, abiotic and biotic stress response pathways, reflecting a greater global stress response with increased phenotype severity. A similar effect was seen in Arabidopsis (Shao et al., 2017), implicating a broader effect than would be conferred by organelle perturbation alone (Fig 2A). Visualizing GO terms associated with enriched pathways using REVIGO (Supek et al., 2011), genes related to stress and calcium signaling were upregulated (Fig S2C), while photosynthesis and chromatin/cell cycle factors were down-regulated, again reflecting global stress behavior (Fig S2D, Table S2).

**Figure 2.**
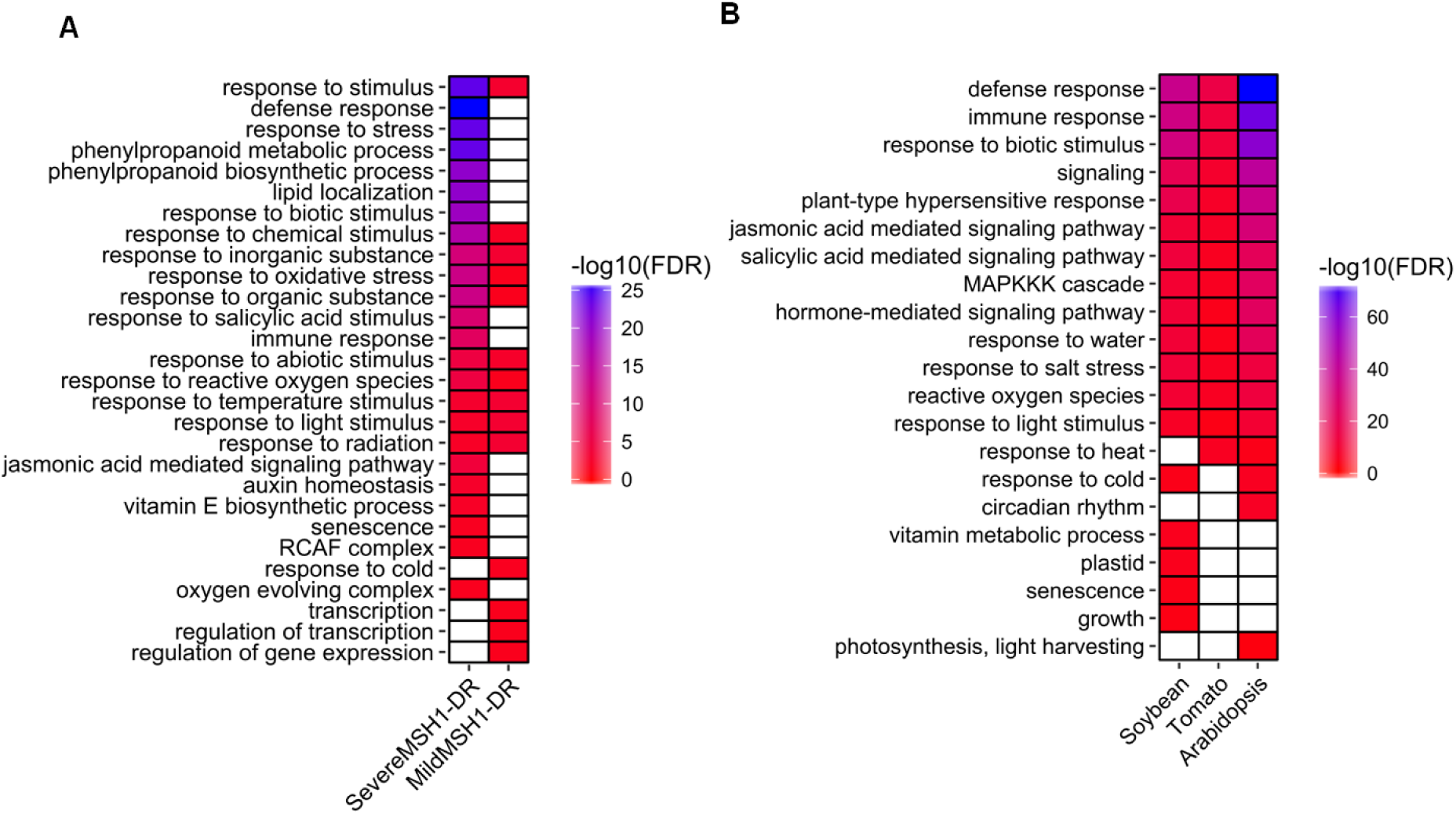
Transcriptome changes in soybean *MSH1*-RNAi lines and cross species comparison of *MSH1*-RNAi gene expression changes. **A)** Heat map differentiating significant GO terms associated with severe and mild *msh1*-RNAi phenotypes (AgriGO, GO term enrichment tool [p-value < 0.05]). **B)** Heat map for conserved and distinct GO terms associated with the *MSH1*-RNAi lines in soybean, tomato, and T-DNA insertion mutant in Arabidopsis. (AgriGO, GO enrichment [p-value < 0.05] was performed on the best Arabidopsis BLAST hit [e-value < e-10] for each soybean and tomato DEG). Heat maps were generated using custom scripts in R.

Cross-species comparison of *msh1*-RNAi soybean transcriptome data with Arabidopsis *MSH1* T-DNA mutant (Shao et al., 2017) and tomato *msh1*-RNAi lines (Yang et al., 2015) showed that while individual genes did not necessarily overlap for differential expression between species, respective GO categories showed high coincidence (Table S3, Fig 2B). Defense, immune response, phytohormone, MAPKKK cascade, biotic and abiotic stress response categories were shared among the three species. We also found differential expression in soybean for orthologs of seven of the 16 signatures belonging to the circadian clock, stress hormone and light-response pathways previously identified through the integration of methylome, RNAseq and network-based enrichment analysis in Arabidopsis *MSH1* memory lines (Yang et al. 2017 submitted; Table S4). Vitamin metabolism and senescence-related genes comprised two categories that were enriched in the soybean *msh1*-RNAi line but not in tomato and Arabidopsis, reflecting a species-specific response to the *msh1*-associated perturbation. The results indicate that *MSH1* suppression confers strikingly similar changes in soybean, tomato, and Arabidopsis in gene expression changes and associated phenotypes.

### Crossing soybean *msh1* memory lines to wild type produces epi-lines with increased variation in adaptive traits

Recent studies have shown that crossing *msh1* memory lines to their isogenic wild type counterpart can influence growth vigor in Arabidopsis, sorghum, and tomato (de la Rosa Santamaria et al., 2014; Virdi et al., 2015; Yang et al., 2015). To investigate the potential of *msh1*-derived vigor in epi-lines of soybean, assess inheritance, and determine the longevity of enhanced growth behavior through self-pollination, we performed reciprocal crosses of *msh1* memory lines with wild type Thorne (Fig S3). Plants in the F_1_ generation were restored to the normal phenotype, ruling out cytoplasmic genetic changes for the *msh1* memory phenotype (de la Rosa Santamaria et al., 2014).

Derived epi-F_2_ lines displayed a broader range of phenotypic variation than wild type for agronomic traits including number of pods (PP) and seeds per plant (SP), seed weight (SW), 100 seed weight (100SW), days to flowering (R1), and days to maturity (R8, Table S5). There was a significant difference in within-genotype variance for number of pods per plant among wild type and the reciprocal F_2_ populations (Fig 3A, Bartlett test, p-value 0.013). The variance estimate for wild type was 103.03, while for WT × T9 F_2_ and T8 × WT F_2_ it was 213.72 and 364.38 respectively. F_2_ populations also differed significantly in flowering time and maturity time, with a small proportion showing higher pod number per plant and delayed maturity (Table S5, Fig 3C).

**Figure 3.**
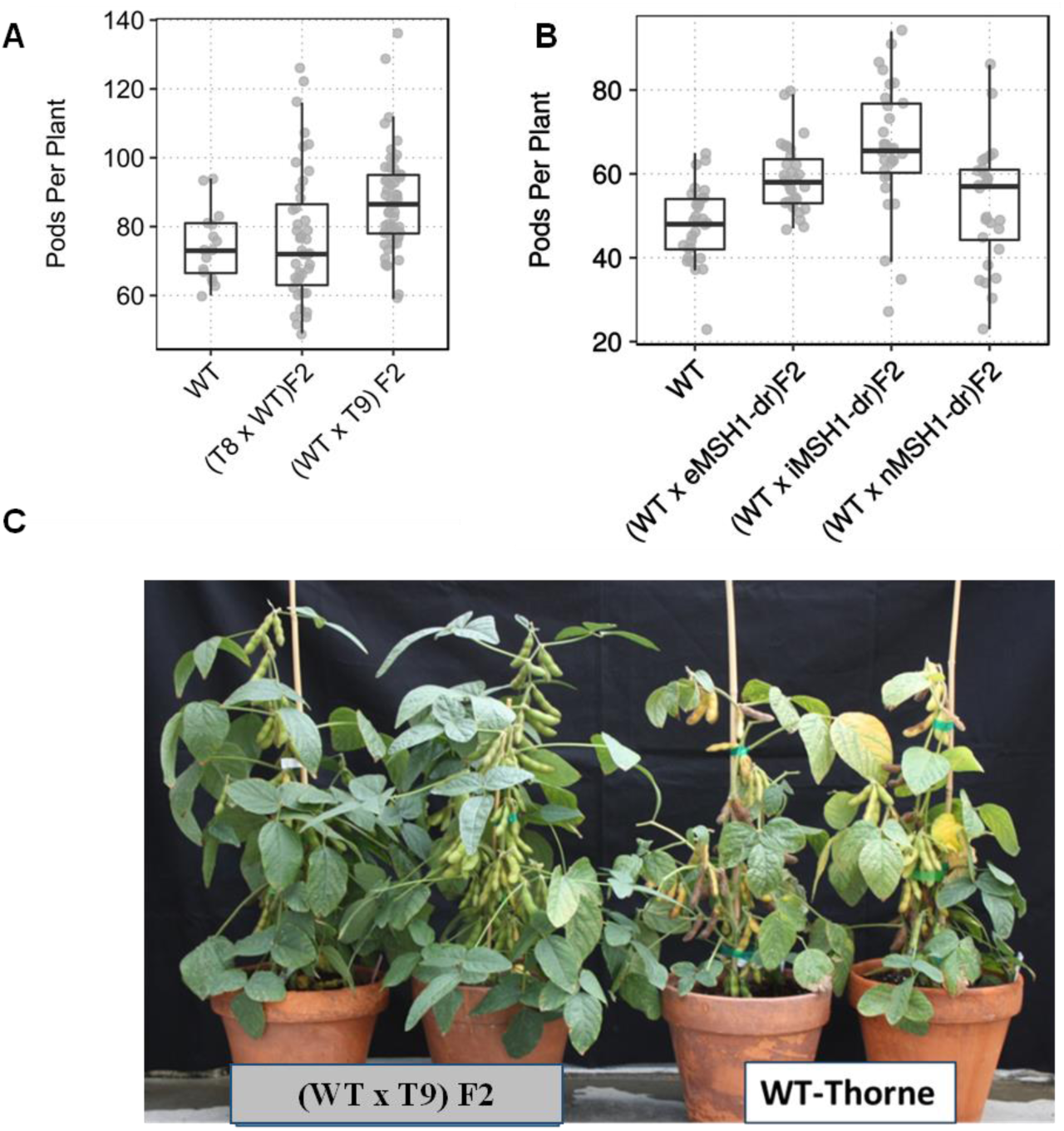
Increased variation for number of pods per plant in different epi-F_2_ populations in the greenhouse. **A)** Enhanced variation for pods per plant in two epi-F_2_ populations compared to wild type grown under greenhouse conditions. **B)** Variation in F_2_ performance for number of pods per plant in the greenhouse for populations derived from a range of *msh1* memory phenotypes (e*MSH1*, i*MSH1*, and n*MSH1*). **C)** WT x T9 epi-F_2_ lines P-37 and P34 showing increase in number of pods per plant and delayed maturity compared to wild type.

We subsequently developed epi-F_2_ populations by crossing wild type with three different phenotypic classes of non-transgenic memory lines, extreme, intermediate and normal phenotype (Fig S1A, B), as pollen donors. Similar to the previous reciprocal crosses, there were significant differences in variance between wild type and the three epi-F_2_ populations for number of pods per plant (Fig 3B, Bartlett test, p-value 0.0011). Increased variance for the measured traits was also observed among the three epi-F_2_ populations. For example, epi-F_2_ WT × eMSH1 showed lower variance than wild type for pods per plant and plant height, but higher variance for days to flowering (Table S6). Epi-F_2_ WT x× iMSH1 had higher variance for pods per plant, and days to flowering, while epi-F_2_ WT × nMSH1 showed higher variance than wild type for all three measured traits (Table S6). These results suggest that *MSH1* epi-populations represent different conditions, perhaps impacting the strategy for selection.

To investigate variation among derived epi-lines and wild type under standard field conditions, we tested 30 F_2:4_ lines from each of the three populations, including 30 wild type sublines as shown in Supplemental Figure S1B. These 120 lines were grown as random complete blocks (RCBD) in four Nebraska locations, Lincoln (SC), Clay Center (CC), Phillips (PH), and Mead (MD), with three replications per location for a total of 12 replications of two-row, ten-foot plots, with rows 3 m long and spaced 0.76 m apart. Data were collected on days to maturity, plant height, protein and oil concentration, and total yield (Table 1).

**Table 1.** Summary of phenotypic data analysis for total yield, maturity, plant height, protein concentration, and oil concentration in wild type and epi-F_2:4_ lines.

| Population* | Yield^ (kg/ha) |  |  | Maturity<sup>#</sup> (DAP) |  |  | Plant Height<sup>§</sup> (cm) |  |  | Protein concentration<sup>%</sup> (percent) |  |  | Oil concentration<sup>%</sup> (percent) |  |  |
| --- | --- | --- | --- | --- | --- | --- | --- | --- | --- | --- | --- | --- | --- | --- | --- |
|  | Mean | SD | Range | Mean | SD | Range | Mean | SD | Range | Mean | SD | Range | Mean | SD | Range |
| **Wild type (Thorne)** | 4374.5 | 361.4 | 3267-5220 | 117.83 | 4.98 | 108-124 | 87.7 | 2.2 | 81-92 | 36.7 | 0.5 | 35.3-37.9 | 19.1 | 0.27 | 18.5-20.0 |
| **F2:4 eMSH1** | 4336.2 | 379.2 | 3049-5143 | 117.5 | 6.79 | 107-127 | 87.1 | 2.12 | 82-91 | 36.7 | 0.57 | 35.0-38.1 | 19 | 0.25 | 18.3-19.7 |
| **F2:4 iMSH1** | 4408.6 | 373.5 | 3471-5303 | 118.64 | 6.04 | 107-126 | 88.5 | 2.39 | 83-93 | 36.9 | 0.48 | 35.6-38.3 | 18.8 | 0.2 | 18.2-19.4 |
| **F2:4 nMSH1** | 4390.9 | 378.5 | 3377-5447 | 116.84 | 4.88 | 108-124 | 86.8 | 2.33 | 82-93 | 36.8 | 0.51 | 35.3-38.1 | 18.9 | 0.24 | 18.3-19.5 |

\* Different epi-population (F2:4 eMSH1, F2:4 iMSH1, and F2:4 nMSH1) were developed from crossing wild type Thorne with *msh1* memory lines varying in phenotypic severity (Supplemental Figure 1B).
^ Yield was measured as total machine harvestable seed weight normalized to 13% moisture.

Similar to greenhouse results for epi-F_2_, we observed differences in variance components for total yield. Epi-F_2:4_ nMSH1 showed ten times higher variance than wild type for total yield, while epi-F_2:4_ eMSH1 showed variance similar to wild type (Table S7). We recorded single plant measurements for pod number per plant, number of branches, and plant height, from ten randomly selected epi-lines in each population along with ten wild type sub-lines from the multilocation field trial. Data were collected from five randomly selected plants from a plot, with two replicates in two locations, Mead and Clay Center. These two locations represent different agro-ecological zones in Nebraska with contrasting soil types. From ANOVA tests, we saw no significant variation among strains or plants within strains for number of branches. For plant height, we saw significant variation among strains in F_2:4_ iMSH1 (p-value = 0.0096) and F_2:4_ nMSH1 (p-value = 0.0075). Epi-F_2:4_ iMSH1 also showed significant variation among strains for pods per plant (p-value = 0.03), while wild type showed significant difference among plants within strains (p-value = 0.007, Table S8). These observations again indicate that epi-lines may differ significantly in their *msh1* effects.

### Selected *MSH1* epi-lines show increased yield compared to wild type in multi-year field trials

To evaluate field performance of *MSH1* epi-lines, F_2:4_ lines were derived from an upper 6% selection for number of pods per plant in the F_2_ generation. Thirty plants each from the selected F_2:3_ lines were grown in the greenhouse and the upper half of these 30 plants, based on number of pods per plant was bulked to form the ‘top 50% selection (S)’. In addition, equal numbers of seeds from all thirty plants were bulked to form the F_2:4_ ‘bulk composite’ (Figure S3). These F_2:4_ lines and wild type were grown as ten-foot two-row plots in the field at Havelock farm in Lincoln, Nebraska, during 2014.

Wild type Thorne showed a mean yield of 4284.65 kg/ha, whereas bulk epi F_2:4_ line yields ranged from 4419.82 kg/ha to 4834.89 kg/ha and top 50% selection epi-F_2:4_ line yields ranged from 4758.33 kg/ha to 5016.7 kg/ha. F_2:4_ R10S yielded significantly better than wild type (Welch two-sample t-test, p-value 0.02, Fig 4A) with a 95% confidence interval for yield gain between 283.3 and 1180.8 kg/ha. As a population, T8 × WT F_4_ yielded 4618.38 kg/ha and WT × T9 F_4_ yielded 4657.85 kg/ha compared to wild type, which yielded 4284.65 kg/ha.

**Figure 4.**
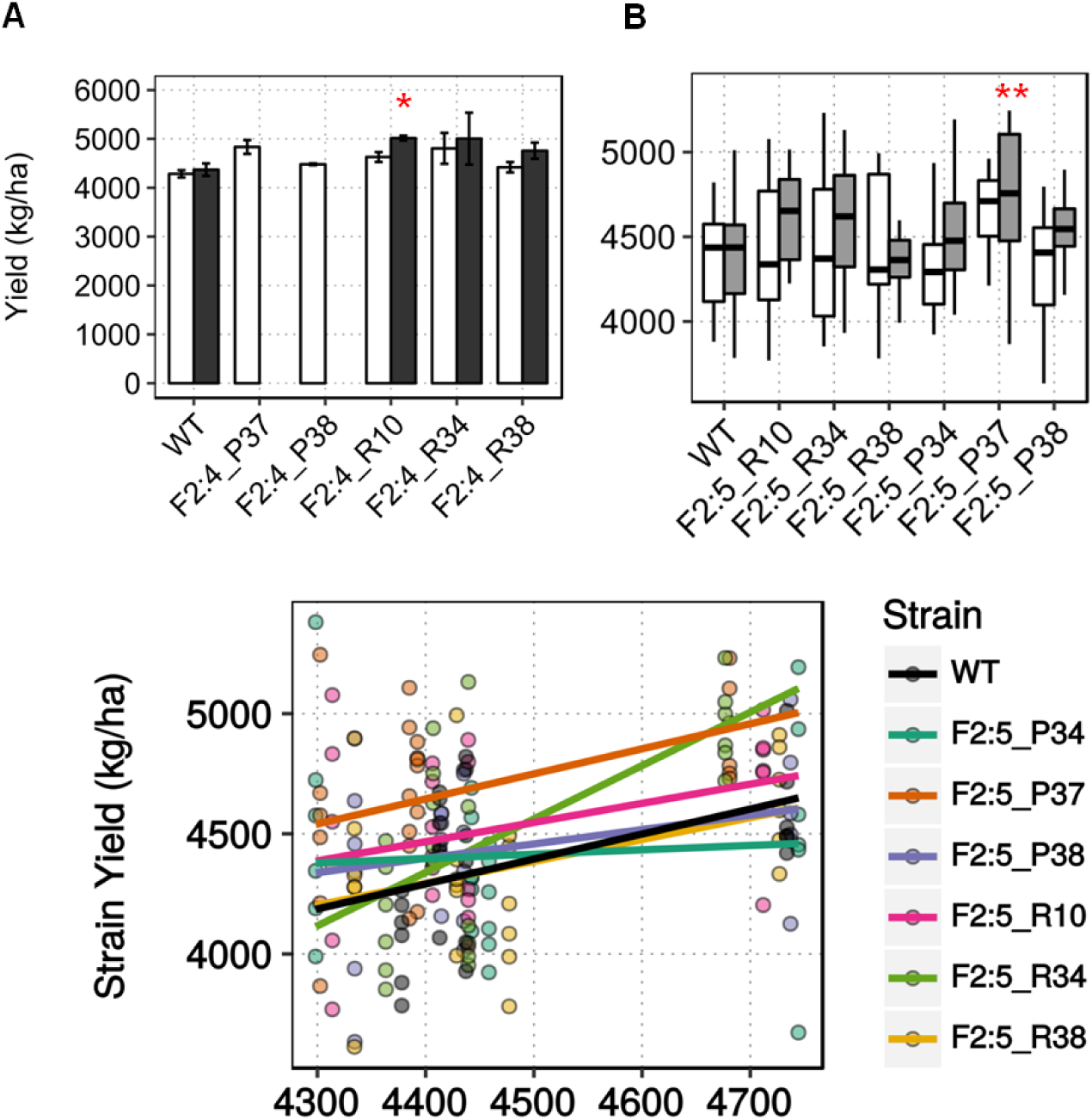
MSH1-derived enhanced growth in field trials. **A)** Enhanced growth measured as total seed weight in kg/ha normalized to 13% moisture for selected epi-F_2_:4 lines in field experiments (n=2). Asterisks represent statistical significance based on Welch two-sample t-test (p-value 0.02) **B)** Mean yield data from pooled locations showing enhanced yield in P37 F_2_:5 epi-line compared to wild type (yield data pooled from three replicates each from four locations). Asterisks denote statistical significance based on t-test (p-value 0.00931) **C)** Reaction norm plots showing superior yield performance of F_2_:5 P37 across environmental index for yield in kg/ha.

Derived F_2:5_ epi-lines (Fig S3) were grown in four different Nebraska locations in the summer of 2015, Lincoln (SC), Clay Center (CC), Phillips (PH) and Mead (MD), with three replications at each site. Mean yield data pooled across locations showed that grain yield for F_2:5_ P37 was significantly higher than wild type by 301.8 kg/ha (t-test, p-value 0.00931, Fig 4B), an increase of seven percent. Except for F_2:5_ R38, all lines showed increased grain yield from 27 kg/ha to 301.8 kg/ha. Regression over an environmental index to visualize any epitype-by-environment (e × E) interactions showed F_2:5_ P34 to have a higher slope than wild type, but not significant by ANOVA. F_2:5_ P37 showed consistently higher yield than wild type across all environmental indices (Fig 4C).

To confirm that there was no penalty for enhanced seed yield in seed quality parameters, we measured seed protein concentration and oil concentration. There was no significant difference in seed protein concentration and 100 seed weight, but epi-lines derived from T8 × WT crosses showed lower oil concentration compared to wild type. F_2:5_ lines from this population also showed earlier maturity compared to wild type (p-value 0.0164). Lodging score did not show variation among the lines tested (Table S9).

F_2:6_ lines, developed from a greenhouse seed increase of 2014-grown epi-F_2:4_, showed no significant difference in mean yield compared to wild type (Fig S4), indicating that the enhanced growth effects taper back to wild type levels by F_6_. Consequently, these experiments demonstrate strongest yield enhancement at F_2:4_ and F_2:5_ generations, with the growth performance returning to wild type levels by F_2:6_, similar to the reported dissipation of epigenetic effects over generations in *ddm1* epiRILs (Cortijo et al., 2014; Roux et al., 2011).

### Progenies of wild type scion grafted on *MSH1*-RNAi show increased yield in field trials

We tested whether enhanced growth could also be observed from *msh1*-grafted progenies in soybean, drawing on previous reports in Arabidopsis and tomato (Virdi et al., 2015; Yang et al., 2015). For this experiment, we grafted three different phenotypic classes of *msh1*-RNAi rootstocks with wild type Thorne scions (Fig S5A), collected seeds from the graft plants, and self-pollinated them for one generation before planting in the 2015 multi-location field trial. Results showed significant yield increase in S2 grafted progenies over wild type (Fig S5B). The type of *MSH1*-RNAi phenotype used as rootstock appeared to make a difference, with WT / n*MSH1*-RNAi lines showing significantly higher yield compared to WT / WT graft (t-test, p-value 0.040) or WT (t-test, p-value 0.019), whereas the WT / i*MSH1* S2 line was marginally better than WT (t-test, p-value 0.052), and WT / e*MSH1* was not significantly different from wild type (Fig S5B). These results further support the non-genetic nature of enhanced growth and the involvement of mobile signals in the process.

### *MSH1*-derived epi-lines are more stable across environments

Epi-F_2:4_ populations developed by crossing wild type with the three different phenotypic classes (e*MSH1*, i*MSH1*, and n*MSH1*) of non-transgenic memory lines were grown together with thirty wild type sub-lines in four locations. The experiment involved a total of 12 replications of two-row, ten-foot plots.

We performed ANOVA tests for interaction between strain and location within populations. As expected, wild type showed strain x location interaction (t-test, p-value 0.0142), but the epi-F_2:4_ populations showed no significant interaction (Fig 5A). To understand this outcome, we plotted the strain means across locations, showing more cross-over interaction for wild type lines, particularly between SC and PH locations (Fig 5B). PH is a higher yielding site, with a mean yield of 4639.9 kg/ha, compared to SC, with a mean yield of 4403.04 kg/ha. Most lines from the epi-population showed an increase in yield from SC to PH, while many of the wild type sub-lines declined. There was also a higher spread of values for wild type sub-lines at the MD location, which may be driving the interaction effects. Epi-lines generally demonstrated higher yield consistency, with F_2:4_ i*MSH1* lines showing tighter grouping in both AMMI plots (Fig S6) and reaction norms (Fig 5B), and performing well in good environments as shown by performance in PH.

**Figure 5.**
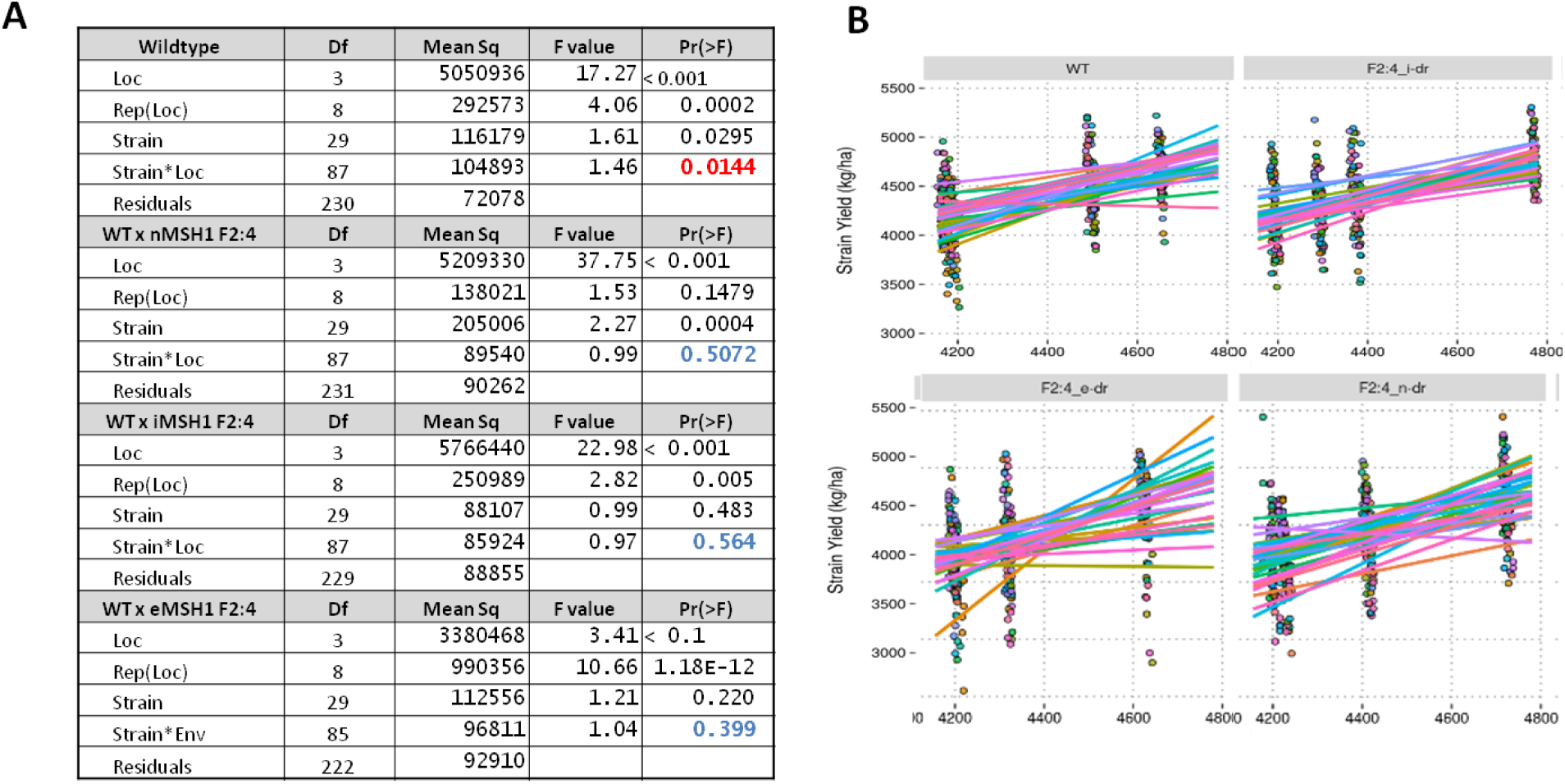
Reduced epitype-by-environment interaction in epi-lines. **A)** Test of significant epitype x environment interaction in wild type sub-lines by ANOVA. **B)** Reaction norm plots based on environmental index for wild type and three different *MSH1* epi-populations with higher cross-over interaction in wild type.

From the variance component estimation, we see that wild type had more than eight fold higher G x E variance estimate than epi-F_2:4_ populations for total yield (Table S7). There was no significant difference in G X E variance component for other traits like maturity date, height, and protein and oil concentrations. From the analysis of single plant measurements for among-strain variation, wild type did not show any significant difference while epi-lines, particularly from F_2:4_ iMSH1, showed significant variation in plant height (p-value 0.0096) and number of pods per plant (p-value 0.03), while F_2:4_ nMSH1 showed significant variation among strains for plant height (p-value 0.007, Table S8). This inherent variation partly explains the buffering capacity for these epi-lines in different environments, leading to reduced e x E interaction. These results imply that *MSH1*-derived vigor and phenotypic plasticity can provide higher yield stability across different environments, although more extensive testing would be necessary to quantify this effect.

### Putative expression signatures in *MSH1*-derived, enhanced growth epi-lines

To investigate biological processes underlying the *MSH1*-derived enhanced yield phenotypes in epi-lines, we performed RNAseq analysis with the two epi-lines R10 and P37 in F_2:4_, F_2:5_, and F_2:6_ generations and their respective wild type controls. These epi-lines showed increased yield in F_2:4_ and F_2:5_ generations, while this enhancement diminished by F_2:6_. We utilized this gradual reversion phenomenon to identify signatures of enhanced growth and their change across generations.

To eliminate the possibility of seed contamination in the epi-lines, we analyzed the RNAseq data with the genome analysis toolkit (GATK) pipeline to identify SNPs from the alignment files. Plotting SNPs across the lines showed no haplotype blocks co-segregating with the enhanced yield lines (Fig S7A, B). When total numbers of SNPs were considered, variation between different epi-lines and wild type was equal to variation within the wild type lines. These data rule out the possibility of seed contamination and are consistent with our hypothesis of epigenetic regulation in *MSH1*-derived epi-lines in the absence of genetic changes.

RNAseq results show R10 F_2:4_ with the greater mean yield gain, to display the highest number of DEGs compared to wild type, with 3048 DEGs, 1259 up-regulated and 1789 down-regulated. R10 F_2:5_ and R10 F_2:6_ showed 955 and 887 DEGs, respectively (Table S10, Fig 6A). We detected 682 DEGs in common between the two epi-lines R10 F_2:4_ and P37 F_2:4_, accounting for 65% of DEGs in P37 F_2:4_ (Fig S8A). GO enrichment (SoyBase) and REVIGO analysis from these DEGs showed up-regulation of stress response pathways (innate immune response, defense, abscisic acid signaling pathway) and down-regulation of metabolism (protein phosphorylation, cellular response to phosphate and magnesium starvation, phosphate ion homeostasis, and galactolipid biosynthesis) (Fig S8B). Several genes related to plastid function and development (Plastid organization, PS II assembly, bilateral symmetry, adaxial/abaxial pattern specificity, response to far-red light, and signal transduction) were differentially expressed only in R10 F_2:4_. Since R10 F_2:4_ was derived from crosses with *msh1* memory line as female parent, these changes are likely remnants of the *msh1* memory effect.

**Figure 6.**
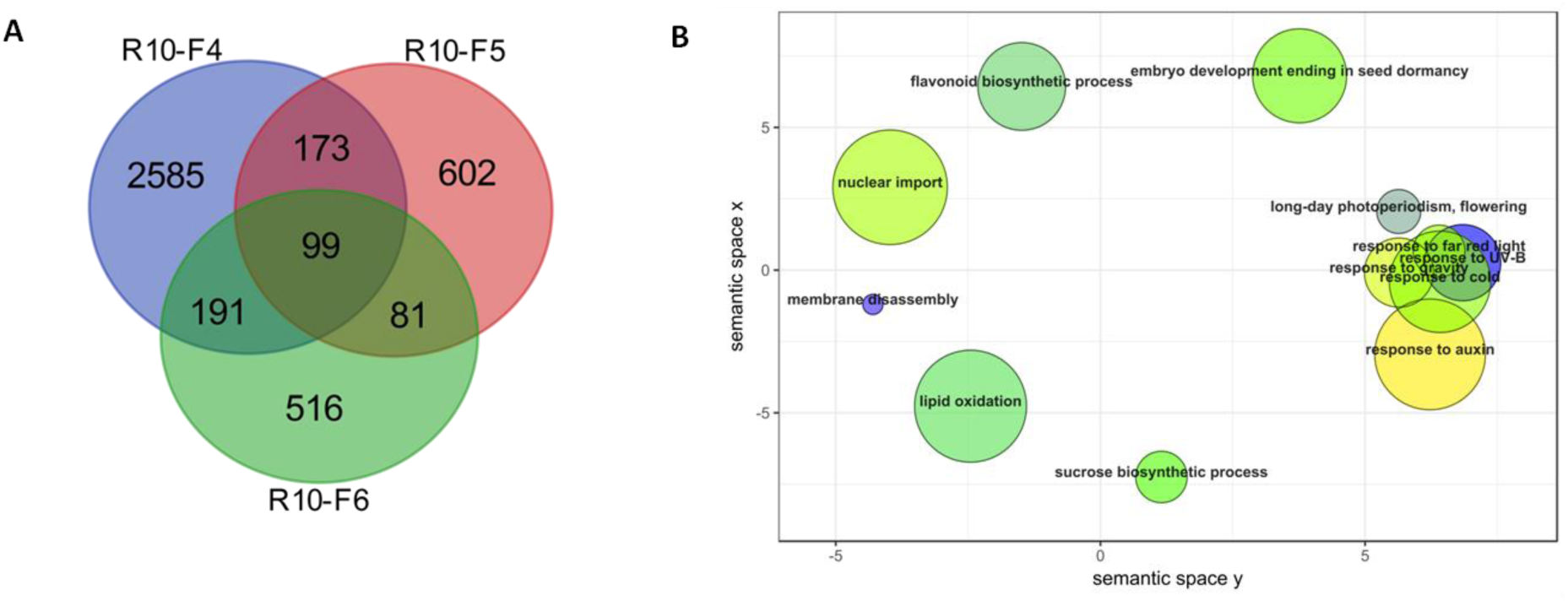
Gene expression changes and associated pathways in *msh1*-derived epi-lines with increased yield. **A)** Venn diagram showing overlap of DEGs in enhanced growth epi-line R-10. **B)** REVIGO plot showing non-redundant GO terms associated with DEGs in epi-line R-10, enhanced growth epi-F_2_:4 compared to epi-F_2_:6, which showed yield similar to wild type. GO terms (p-value < 0.05) obtained from SoyBase were used in REVIGO tool from AgriGO with modified R script for plotting.

To identify signature gene expression changes underlying the enhanced growth effect in epi-F_2:4_ lines and the return to wild type levels by epi-F_2:6_, we compared gene expression changes between F_2:4_ and F_2:6_ within the same lineage. To ensure direct comparison, we omitted genes that were differentially expressed in epi-F_2:6_ vs WT_F6_ and WT_F4_ vs WT_F6_ comparisons. This resulted in a filtered set of 545 DEGs in R10 and 454 DEGs in P37.

Auxin response genes were consistently modulated in both R10 and P37 epi-lines. In R10 F_2:4_ vs F_2:6_ comparisons, we detected changes predominantly in sucrose biosynthesis as well as gravitropism and auxin stimulus response pathways (Fig 6B, Table 2), whereas in P37 F_2:4_ vs F_2:6_ comparisons, genes related to auxin response and protein phosphorylation were enriched (Table 2). A total of 40 DEGs (ca 8%) were common between the two epi-lines. These genes represented auxin response, cell wall and cell cycle, and stress related genes (Table 3). The 40 genes were not necessarily modulated in the same direction in the two epi-lines, perhaps emphasizing the role of circadian regulators in modulating the expression of these genes (Yang et al. 2017, submitted).

**Table 2.** Enriched GO terms associated with *MSH1* derived enhanced growth in R10 and P37 epi-lines.

| Type | GO_id | GO_count | Expressed | Expected | P_adj | GO_desc |
| --- | --- | --- | --- | --- | --- | --- |
| **R-10F4 vs R-10F6\*** | GO:0005986 | 35 | 6 | 0.5 | 0.0049 | sucrose biosynthetic process |
|  | GO:0009629 | 23 | 5 | 0.3 | 0.0083 | response to gravity |
|  | GO:0009733 | 1020 | 31 | 14.3 | 0.0408 | response to auxin stimulus |
| **P-37F4 vs P-37F6<sup>#</sup>** | GO:0009733 | 1020 | 26 | 10.4 | 0.0127 | response to auxin stimulus |
|  | GO:0006468 | 2386 | 7 | 24.4 | 0.038 | protein phosphorylation |

\* Represents DEGs between enhanced growth epi-line R-10 F<sub>2:4</sub> (derived from epi-population with *msh1* memory line as female parent) compared to R-10 F<sub>2:6</sub> line with yield similar to wild type.
<sup>#</sup> Represents DEGs between enhanced growth epi-line P-37 F<sub>2:4</sub> (derived from epi-population with *msh1* memory line as pollen donor) compared to P-37 F<sub>2:6</sub> line with yield similar to wild type.

**Table 3:** Common DEGs in two enhanced growth epi-F_2:4_ lines, R-10 and P-37 compared to its respective epi-F_2:6_.

| **Auxin biosynthesis related** |  |  |  |
| --- | --- | --- | --- |
| Glyma.03G158700 | AT4G14550 | IAA14 | indole-3-acetic acid inducible 14 |
| Glyma.04G006900 | AT5G18060 |  | SAUR-like auxin-responsive protein family |
| Glyma.06G281800 | AT4G38840 |  | SAUR-like auxin-responsive protein family |
| Glyma.06G282000 | AT4G38840 |  | SAUR-like auxin-responsive protein family |
| Glyma.06G282100 | AT4G38840 |  | SAUR-like auxin-responsive protein family |
| Glyma.06G282600 | AT5G18020 |  | SAUR-like auxin-responsive protein family |
| Glyma.06G282700 | AT4G38840 |  | SAUR-like auxin-responsive protein family |
| Glyma.07G034200 | AT3G15540 | IAA19 | indole-3-acetic acid inducible 19 |
| Glyma.12G141000 | AT5G54510 | GH3.6 | Auxin-responsive GH3 family protein |
| Glyma.12G226600 | AT3G15210 | ERF4 | ethylene responsive element binding factor 4 |
| **Cell cycle/growth related** |  |  |  |
| Glyma.01G035600 | AT1G70210 | CYCD1;1 | CYCLIN D1;1 |
| Glyma.03G171400 | AT5G59970 |  | Histone superfamily protein |
| Glyma.04G166700 | AT1G26550 |  | FKBP-like peptidyl-prolyl cis-trans isomerase family |
| Glyma.05G002500 | AT3G01640 | ATGLCAK | glucuronokinase G |
| Glyma.07G133800 | AT5G02220 | SMR4 | Cyclin dependant kinase inhibitor |
| Glyma.08G277700 | AT5G13420 |  | Aldolase-type TIM barrel family protein |
| Glyma.08G287500 | AT1G70370 | PG2 | polygalacturonase 2 |
| Glyma.09G073600 | AT3G43190 | SUS4 | sucrose synthase 4 |
| Glyma.09G189700 | AT5G53250 | AGP22 | arabinogalactan protein 22 |
| Glyma.11G011000 | AT3G04500 |  | RNA-binding (RRM/RBD/RNP motifs) family protein |
| Glyma.14G219100 | AT1G75750 | GASA1 | GAST1 protein homolog 1 |
| Glyma.15G093700 | AT4G18340 |  | Glycosyl hydrolase superfamily protein |
| Glyma.15G109800 | AT4G04470 | PMP22 | Peroxisomal membrane 22 kDa (Mpv17/PMP22) family |
| Glyma.17G140000 | AT4G12510 |  | Seed storage 2S albumin superfamily protein |
| Glyma.18G206000 | AT2G38310 | PYL4 | PYR1-like 4, ABA Signalling |
| Glyma.19G069200 | AT1G07430 | HAI2 | highly ABA-induced PP2C gene 2 |
| Glyma.19G206300 | AT1G03470 |  | Kinase interacting (KIP1-like) family protein |
| **Stress response related** |  |  |  |
| Glyma.01G060300 | AT1G13740 | AFP2 | ABI five binding protein 2 |
| Glyma.04G003300 | AT2G47140 |  | NAD(P)-binding Rossmann-fold superfamily protein |
| Glyma.04G003700 | AT4G38580 | ATFP6 | farnesylated protein 6 |
| Glyma.05G149400 | AT4G10340 | LHCB5 | light harvesting complex of photosystem II 5 |
| Glyma.05G153200 | AT5G13930 | CHS | Chalcone and stilbene synthase family protein |
| Glyma.11G179100 | AT1G08440 |  | Aluminium activated malate transporter family protein |
| Glyma.14G093100 | AT3G09390 | MT2A | metallothionein 2A |
| Glyma.15G251500 | AT1G78380 | GST8 | glutathione S-transferase TAU 19 |
| Glyma.16G121900 | AT2G17730 | NIP2 | NEP-interacting protein 2 |
| Glyma.16G215800 | AT5G54300 |  | Chloroplast membrane protein unknown function |
| Glyma.17G140700 | AT4G16520 | ATG8F | Ubiquitin-like superfamily protein, plastid autophagy |
| Glyma.18G111300 | AT2G30860 | GSTF9 | glutathione S-transferase PHI 9 |
| Glyma.20G140400 | AT4G05200 | CRK25 | cysteine-rich RLK (RECEPTOR-like protein kinase) 25 |

Auxin response genes include IAA19, a positive regulator of plant growth (Kohno et al., 2012) required for PIF4-mediated modulation of auxin signaling (Sun et al., 2013). SMALL AUXIN UP RNAs (SAURs) were differentially expressed in both epi-lines. SAUR genes are involved in cell expansion and development, particularly for integrating hormonal and environmental signals that regulate plant growth (Li et al., 2015; Ren and Gray, 2015). These data provide candidate pathways underpinning the growth behavior in *MSH1* epi-lines.

## DISCUSSION

Previous studies have shown the influence of *MSH1* perturbation for altering growth in Arabidopsis, sorghum, and tomato (de la Rosa Santamaria et al., 2014; Virdi et al., 2015; Yang et al., 2015). The present study exploits epigenetic variation induced by *MSH1* perturbation in soybean to develop epi-lines that displayed an increase in seed yield from selected F_4_ and F_5_ families, subsiding by the F_6_ generation, under large-scale field conditions. Epi-lines showed reduced epitype-by-environment interaction, inferring contribution of the *MSH1* effect to buffering across environments. Gene expression profiling within the derived epi-lines uncovered pathways modulated in the enhanced growth F_4_ and F_5_ cycles that returned to wild type levels by F_6_. Effects were particularly pronounced in auxin response pathways, suggesting their possible utility as candidate markers in early selection. Observation of auxin response pathway modulation in tomato epi-lines further strengthens this argument (Yang et al., 2015).

Disruption of *MSH1* in both monocot and dicot plant species produces remarkably similar developmental reprogramming phenotypes that are independent of transgene segregation (Xu et al., 2012). Apart from conditioning a similar phenotypic response, *MSH1* disruption in various plant species show similar transcriptome behavior, including changes in immune and defense, circadian rhythm, phytohormone, and abiotic stress response pathways (Fig 2B). Methylome behavior in *msh1* memory lines of Arabidopsis and tomato are also consistent (Yang et al. 2017, submitted), further reiterating cross-species conservation for the *MSH1* effect.

In Arabidopsis, epigenome disruption through crossing wild type *Col-0* with *met1*-derived epiRILs triggers reprogramming of DNA methylation and changes in gene expression patterns in the F_1_ epi-hybrids (Rigal et al., 2016). Similarly, crossing soybean *msh1* memory lines to isogenic wild type brings together two genetically identical genomes varying in DNA methylation patterns, creating conditions for widespread changes in DNA methylation and gene expression. The increased phenotypic variation in agronomic traits seen in F_2_ populations may be a consequence of segregation of these novel methylation patterns and chromatin changes.

Derived F_2:4_ epi-lines showed significant variation for agronomically important traits like yield and days to maturity. Increasing variation in the population is considered beneficial under challenging conditions (Herman et al., 2014). Similar to bet hedging under different environments, epigenetically variable lines should be favored, since a portion of the individuals are more suited to the prevailing environmental conditions, providing buffering capacity for the population (Herman et al., 2014). Our data, albeit early-stage, support this notion by displaying reduced epitype-by-environment interaction than was observed in isogenic wild type across environments.

All six lines selected from top performing F_2_ plants showed a reduction of enhanced growth by F_2:6_, further confirming the epigenetic nature of *MSH1*-derived growth changes, with similar dissipation patterns described previously in Arabidopsis *ddm1* epiRILs (Cortijo et al., 2014; Roux et al., 2011). A recent study has suggested that stability and switching of acquired epigenetic states are influenced by DNA sequence composition and repetitiveness (Catoni et al., 2017). It is also speculated that methylation variation not linked to a causal genetic variant tends to be less stable than when directly linked to genetic change (Schmitz et al., 2013). We deployed a strict top 6% selection in the F_2_ generation from each population for further evaluation. We assume that a more relaxed selection from these populations might show sustained enhanced growth for extended generations beyond F_2:6_.

GO enrichment analysis of DEGs in derived epi-lines with increased yield showed changes in genes associated with photosynthesis, plastid organization, defense, immune response, light response, and phytohormones. These pathways are also differentially modulated in *msh1* mutants (Shao et al., 2017), and similar gene expression changes in stress and phytohormone pathways are seen in *MSH1*-derived epi-F_3_ lines of tomato (Yang et al., 2015).

Soybean epi-line R10 F_2:4_ (*msh1* memory line as female parent) showed greater correspondence with the gene expression patterns of *msh1* mutants than did epi-line P-37 F_2:4_ (wild type as female parent). These observations suggest that the *msh1* mutant profile represents both organellar and epigenetic contributions to a global gene expression repatterning, and we are seeking to further dissect this phenomenon.

Immune and defense response genes were consistently upregulated in the two soybean epi-lines, in contrast to their repression in Arabidopsis F_1_ plants from ecotype hybrids (Groszmann et al., 2015; Miller et al., 2015), perhaps reflecting a fundamental difference between *MSH1* derived enhanced growth and heterosis. Comparison of gene expression changes in F_2:4_ vs. F_2:6_ within a single lineage offers a unique system to understand the pathways associated with enhanced growth in the *MSH1* system. By this analysis, auxin response genes emerge in both epi-lines tested to date, consistent with previous reports from Arabidopsis ecotype hybrids (Groszmann et al., 2015; Wang et al., 2017) and with previous studies in *msh1*-derived tomato epi-F_3_ lines (Yang et al., 2015).

SAUR genes are implicated in regulating plant growth through sensing hormone and environmental cues (Li et al., 2015; Ren and Gray, 2015). These genes encode small proteins unique to plants that are found in tandem arrays or as segmental duplications of closely related genes (McClure and Guilfoyle, 1987) so that assigning a function to each SAUR gene is challenging. Recent evidence suggests an emerging relationship between phytohormones and epigenetic changes like histone modification, chromatin remodeling, and DNA methylation repatterning. Thus, coordinated changes in epigenomes may be one of the outcomes of plant hormone cross talk (Yamamuro et al., 2016).

Sucrose biosynthetic pathway genes were also differentially expressed in R10 F_2:4_ relative to R10 F_2:6_. Starch metabolism changes in Arabidopsis serve as a means to enhance biomass and oil-seed production while maintaining oil quality parameters (Liu et al., 2015). Sucrose synthase (SUS) enzymes play an important role in storage-reserve accumulation in Arabidopsis (Fallahi et al., 2008) and, similarly, fructokinases (FRKs) are important for storage-reserve accumulation and embryo carbon catabolism (Stein et al., 2016). Whether these molecular signatures, both phytohormonal and metabolic, can be exploited in early-generation selection to predict superior epi-lines needs to be investigated further.

We provide evidence that novel epigenetic variation induced by *MSH1* suppression, following crossing and F_2_ segregation, can be inherited for at least three generations and bred for crop improvement with few rounds of selection to enhance and stabilize crop yield. It is unclear whether enhanced phenotypic plasticity will extend beyond this generational timeframe. This is a particularly intriguing question as relates to grafting, where no crossing is involved. These findings have interesting implications for plant breeding, epigenetics, and transgenerational inheritance of non-genetic variation within plant genomes. The observed outcomes portend the utility of induced epigenetic variation within elite inbred lines, exploiting this variation to further enhance and stabilize agronomically important traits. One limitation of our study was that all the lines tested in the multi-location and multi-year experiments were derived from only five different crosses and a similarly limited number of graft events, making it difficult to assess the frequency and effect of *msh1* memory and *MSH1* suppression phenotypes on crossing and large-scale grafting outcomes. More work is now needed on molecular signatures of the ideal *msh1* suppression and memory lineages that will predict downstream performance and durability of the epigenetic effect.

## MATERIALS AND METHODS

### RNAi Constructs and Transformation

A 557-bp segment encoding amino acids 945 to 1131, which represents the region following the ATPase domain (V) and spanning to the end of the GIY-YIG homing endonuclease domain (VI) of the soybean *MSH1* gene was PCR-amplified using primers Soy-MSF4 (5’-ATCAGTTGGTTTATGCTAAGGAAATGCT-3’) and Soy-3Rbam (5’-TATGTATACAGGTTGGAAGTGCCAAAATTCCTATG-3’). The PCR-amplified fragment was cloned in forward and reverse orientation flanking the second intron of the Arabidopsis small nuclear riboprotein (At4g02840) in the pUCRNAi vector provided by Dr. H. Cerutti (University of Nebraska-Lincoln) and later transferred into pPTN200 (pPZP family binary vector), which carries the *BAR* gene with nopaline synthase promoter and 3’UTR terminator. The hairpin sequences were placed under the control of 35S Cauliflower Mosaic Virus (CaMV) promoter with a duplicated enhancer and terminated by its 3' UTR. The final vector CIPB-7 was used to transform soybean by the cotyledonary node method of *Agrobacterium-mediated* transformation (Xing et al., 2000; Zhang et al., 1999), and the herbicide Basta was used for selection of transformants.

### Plant Material and Growth Conditions

For greenhouse studies, seeds were sown into moist peat pots containing standard potting mix, and transferred to 8” pots after two weeks. Plants were grown under 16-hr light/dark cycle at 28^0^C. Days-to-flowering (R1) was measured as number of days from sowing to one open flower at any node on the stem. Days-to-maturity (R8) was measured as number of days from sowing until 95% of pods were mature and brown. Plant height was taken at R8 developmental stage as distance between the soil surface and the apical meristem of the main stem. All plants were hand-harvested individually, and number of pods was recorded before threshing to obtain number of seeds per plant. Near infrared (NIR) technology was used to determine protein concentration, oil concentration, and moisture content of the seeds. Total seed weight was normalized to 13% moisture level.

Grafting was performed in the greenhouse on *MSH1*-RNAi and *msh1* memory lines. Wild type seedlings at 12-14 days after sowing were used as scion and grafted onto 10-day-old root stocks of wild type control or *MSH1* lines by the wedge grafting technique (Bezdicek et al., 1972; Kiihl et al., 1977) with necessary modifications. Seeds were collected from the grafted scion and 30 plants from each graft was grown for one generation (S1) in the green house and bulk harvested to obtain S2 seeds. Graft-S2 lines were grown as two-row, ten foot plots in multilocation field trials with three reps in each location for a total of 12 replications per graft.

During 2014 summer, twelve epi F_2:4_ lines with wild type were grown as four row plots (3m long and 0.76m apart). All data, including plot yield, were collected on the center two rows of each plot. Emergent seedlings in each plot were counted two weeks after sowing to determine seed density and four epi-lines which had lower than 24 seeds per meter were discarded from further analysis. All lines were grown in a completely randomized design with two replicates. Rows were hand-harvested and threshed on site, and grain yield measured as total seed weight for the plot adjusted to 13% moisture and converted to Kg/ha.

In 2015, a multi-location trial was conducted at four different Nebraska locations: Lincoln, Mead, Clay Center and Phillips. Lines were grown as two-row plots (2.9m long and 0.76m apart) with 24-26 seeds per meter. In separate experiments, 12 F_2:5_ lines and six F_2:6_ lines from the reciprocal cross experiment were grown as random complete blocks (RCBD) with three replications at each location. In another experiment, 30 epi-lines each from three epi-F_2:4_ populations were grown along with 30 wild type sub-lines in RCBD with three replicates in four locations. Grain yield was measured as combined harvestable seed yield adjusted to 13% moisture. Height was recorded as average length of the main stem from soil surface to tip of the plant, expressed as the average of three individual plants in a uniform section of the row. Maturity date was recorded as number of days from planting until R8 stage, and lodging was scored from 1 to 5, with 1 indicating all plants in the plot erect, 3 indicating a plot average of plants at a 45 degree angle, and 5 showing all plants prostate on the ground. Single plant measurements were recorded from ten randomly selected lines in each population in two locations, Mead and Clay Center, with two replicates. In each plot, five randomly selected plants were marked and measurements were taken for pods per plant, number of branches, number of nodes, and height.

### Phenotypic data analysis

For ANOVA analysis of main effects and interactions in 2014 and 2015 field experiments, trait values were first fitted using the “lm” function in R with the linear model *y_ijk_* ~ *line_i_* + *env_j_* + (*line*env*)*_ij_* + (*rep/env*)*_kj_* + *e_ijk_*, where *line_i_* is the main effect of line *i*, *env_j_* is the main effect of environment *j*, (*line*env*)*_ij_* is the interaction between line *i* and environment *j*, (*rep/env*)*_kj_* is the effect of replicate *k* nested within environment *j*, and *e_ijk_* is the residual error; all independent variables were treated as fixed effects. Tests for significant effects and interactions were then performed using the ANOVA function within the “car” R package. In the 2015 multilocation trial, outliers for grain yield were identified based on a threshold of more than 2x the interquartile range below the first quartile or above the third quartile (resulting in 4 observations removed).

For phenotypic analysis within and across multiple environments (2015 multi-location trial), mean trait values and corresponding confidence intervals were estimated for each line using the “lme4” R package with the linear mixed model *y_ijk_* ~ *line_i_* + *env_j_* + *(line*env)_ij_* + *(rep/env)_kj_* + *e_ijk_*, where *line_j_* was treated as a fixed effect and (*rep/env)_kj_* was treated as a random effect. Tests for significant differences in line means were performed using general linear hypothesis tests with the “multcomp” R package, with p-values adjusted using the Benjamini-Hochberg method. After fitting the model, variance components were extracted using “VarCorr()” function in R. For analysis of single plant measurements in the field to look at strain variance and within line variance, data analysis was done using proc glm in SAS.

Joint regression analysis (Finlay and Wilkinson, 1963) was performed to assess individual line performance relative to the grand population performance across environments (i.e. environmental index). Trait data values for each line were regressed over the mean trait performance of all lines within that environment, excluding the line being estimated to avoid bias (Wright, 1976); the resulting slope of each line is an indicator of its response to environmental change compared to the population mean (Lynch and Walsh, 1998). AMMI plots were generated using the “agricolae” R package.

### Microarray, RNA-seq and SNP analysis

RNA preparation and processing for microarray assay has been described previously (Xu et al., 2011). We performed Gene Ontology (GO) analysis by converting the Affy probe ID into Soybean Genome ID (Phytozome) using a custom script in R. AgriGO (Du et al., 2010) analysis was performed on this list of differentially expressed genes. For comparative analysis, the best Arabidopsis BLAST hit for each differentially expressed orthologous gene in *MSH1*-RNAi tomato (Yang et al., 2015) and severe *MSH1*-RNAi soybean was used to generate GO enrichment and plotted as a heat map using custom R scripts.

For RNAseq, leaves from four-week-old plants were harvested and frozen in liquid N2. Three biological replicates for each epi-line, R10 and P37 from F_2:4_, F_2:5_ and F_2:6_ generations were sampled along with three generations of wild type (WT_F4_, WT_F5_, and WT_F6_). RNA was isolated with TRIzol (Invitrogen), followed by RNeasy (Qiagen) column purification. Sequencing was performed by BGI, generating 2x100 bp paired-end reads with a mean of 25.6 million pairs per sample. After trimming bases below a quality score of 20, reads were aligned to the *Glycine max* reference obtained from Phytozome (cv. Williams 82, assembly v2.0) using STAR 2-pass method (Dobin et al., 2013) and allowing a mismatch rate of 0.04*(read length).

This resulted in a mean unique mapping rate of 93.2%, or 97.3% when including multi-mapped reads. From STAR 2-pass alignment files, SNP detection was performed using the Genome Analysis Toolkit (GATK) pipeline. SNP information from all samples was combined to create a total possible SNP list, filtered to only include SNPs supported by an alternate allele frequency of ≥ 0.75 and a read depth of ≥ 10. For every sample, if a SNP was not detected in a given position, it was assumed to be equal to the reference nucleotide. Only positions declared as SNPs in at least two of the 27 samples sequenced were retained as variable sites. Next, every sample was compared against the wild type samples of the other generations as the control, so that the wild type samples could also be evaluated, e.g. WT_F4_, R-10_F2:4_, and P-37_F2:4_ were compared against WT_F5_ and WT_F6_. If a position had a different nucleotide than the wild type samples (only positions with agreement amongst the wild type controls were considered), then it was considered a SNP relative to the wild type Thorne in our material.

All such SNPs were then plotted as depicted in Supplemental Fig S7. Putative SNP haplo-blocks did not co-segregated with higher performance. For differentially expressed genes, reads were mapped to annotated genes (assembly 2, version 1, release 275), then counted with strand-specificity enforced. The Bioconductor package ‘sva’ was used to identify and remove a single surrogate variable related to sequencing lane batch effect. DESeq2 (Love et al., 2014) was used to normalize counts, estimate gene expression, and identify differentially expressed genes (absolute log2 fold-change ≥ 0.5 and a FDR < 0.05). SoyBase (https://soybase.org/) was used for GO enrichment analysis and heat maps generated using custom R scripts.

## ACKNOWLEDGMENTS

We thank the UNL transformation core facility for soybean transformation and Travis Scheuler, Daniel Jaber, Aaron Hoagland, and John Rajeswki for help with field experiments. This work was partially supported by grants from National Science Foundation (IOS1126935) and The Bill and Melinda Gates Foundation (OPP1088661) to S.M.

## AUTHOR CONTRIBUTIONS

Conceptualization: SM and SKKR; Performed experiments: SKKR, YZX, and AS; Data Analysis: SKKR, MSR, GG, and RS; Writing - Original Draft: SKKR; Writing - Review and Editing: SM and GG; All authors read and approved the final manuscript.

---

**Figure S1.**
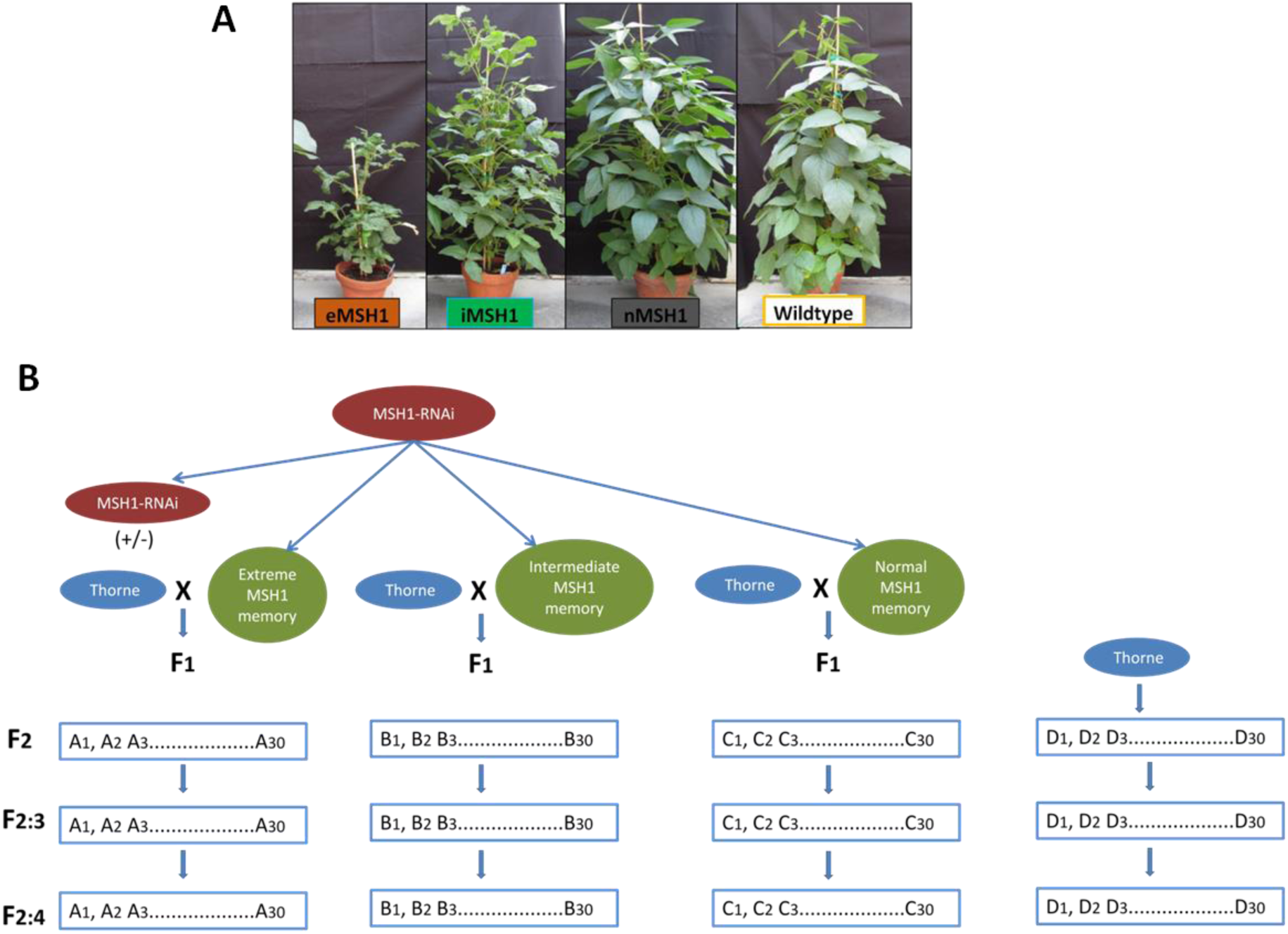
Classification of *MSH1* memory phenotypes into extreme (e*MSH1*), intermediate (i*MSH1*) and normal phenotype (n*MSH1*) **A)** The *msh1* memory lines were classified into categories; *nMSH1* for plants without visible phenotype distinguishing them from wild type, *iMSH1* for plants with intermediate phenotype (leaf, floral alterations and delayed flowering but normal height), e*MSH1* for extreme plants showing dwarfing along with other described phenotypes. **B)** Schematic representation of crossing strategy used to develop epi-populations derived from crossing wild type with three different *msh1* memory lines (e*MSH1*, i*MSH1*, and n*MSH1*)

**Figure S2.**
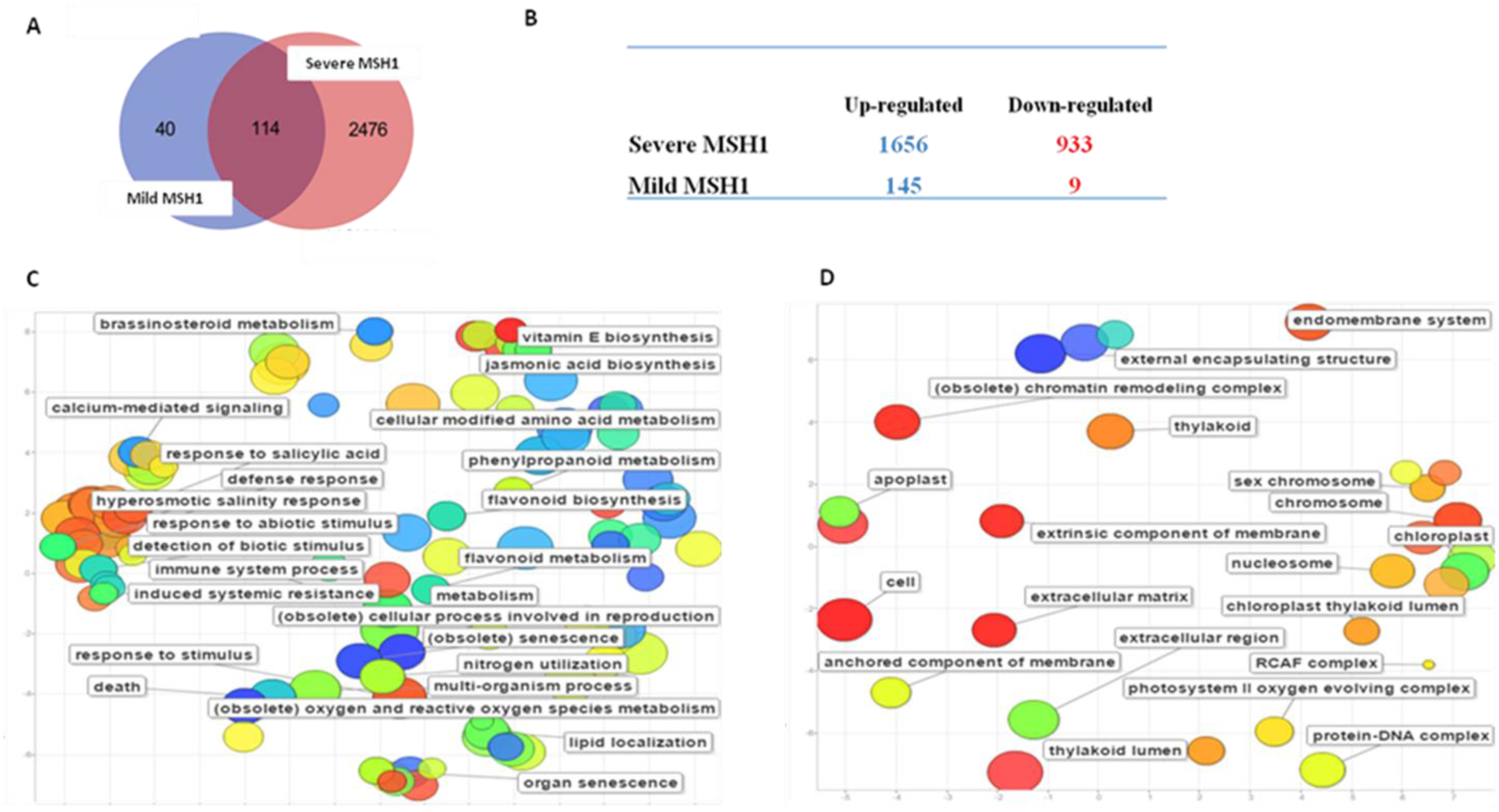
Gene expression changes and ReviGO terms associated with soybean severe *MSH1*-RNAi lines. **A)** Venn diagram showing number of DEGs (p-value < 0.05 and |log2(value)| > 1) in severe and mild *MSH1*-RNAi lines. **B)** Table showing number of up and down-regulated genes in severe and mild *MSH1*-RNAi lines (p-value < 0.05 and |log2(value)| > 1). **C)** REVIGO terms associated with up-regulated and down-regulated genes **(D)** in severe *MSH1*-RNAi. GO terms (p-value < 0.05) obtained from SoyBase were used for REVIGO analysis using default parameters in agriGO. REVIGO summarizes list of GO terms into semantic similarity based scatter plots.

**Figure S3.**
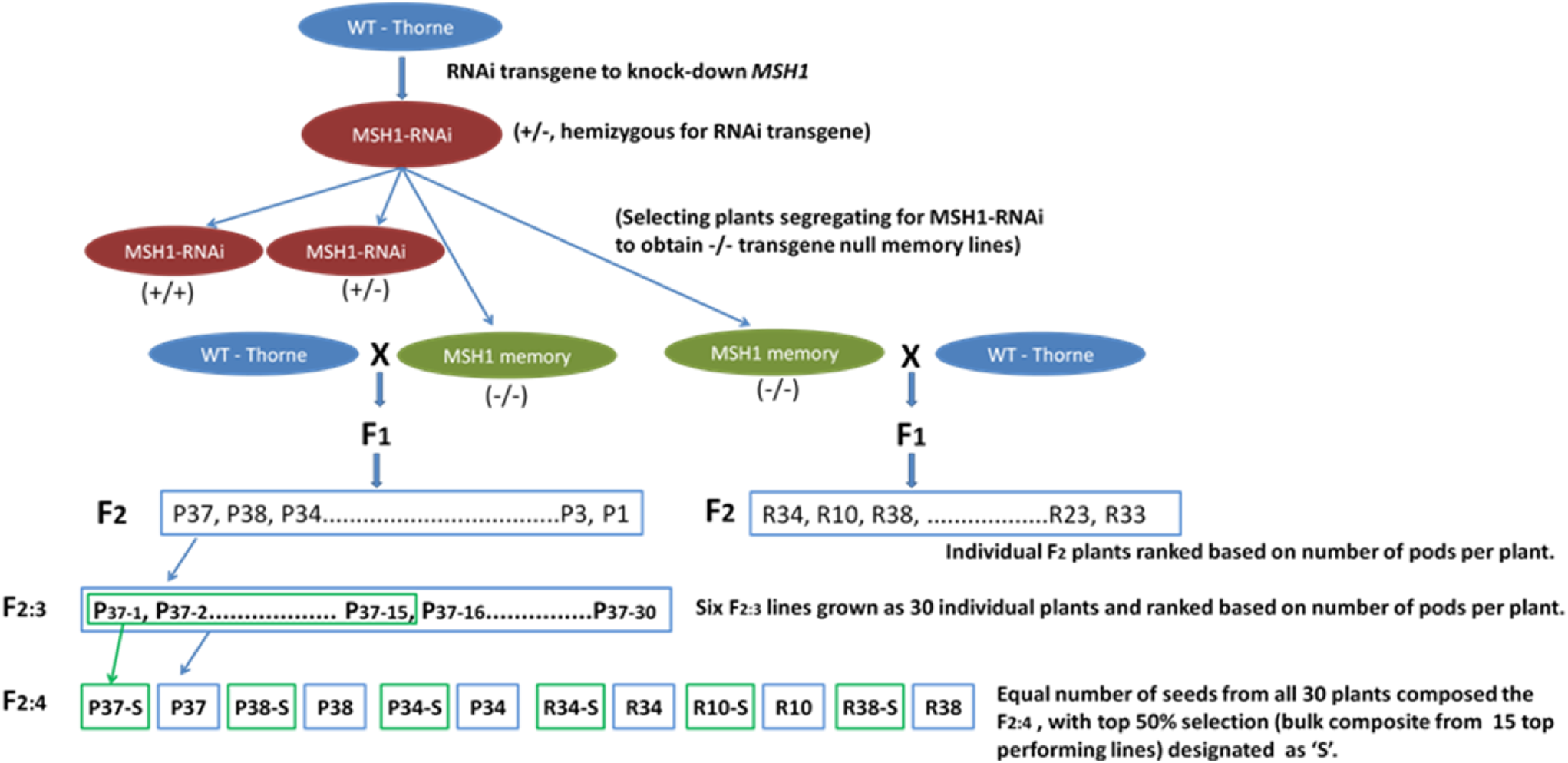
Schematic representation of crossing scheme in *msh1* derived epigenetic breeding. Schematic representation showing *MSH1*-RNAi lines and development of *msh1* memory lines from plants segregating for the transgene. These transgene-null plants (*msh1* memory) with or without phenotype were crossed to wild type in both directions and the F1 self-pollinated to obtain further filial generations. In the F_2_ generation, upper 6% selection was performed on the two populations along with wild type based on number of pods per plant. Another round of selection for upper 50% was done in F_2_:3 lines and forwarded to field experiments.

**Figure S4.**
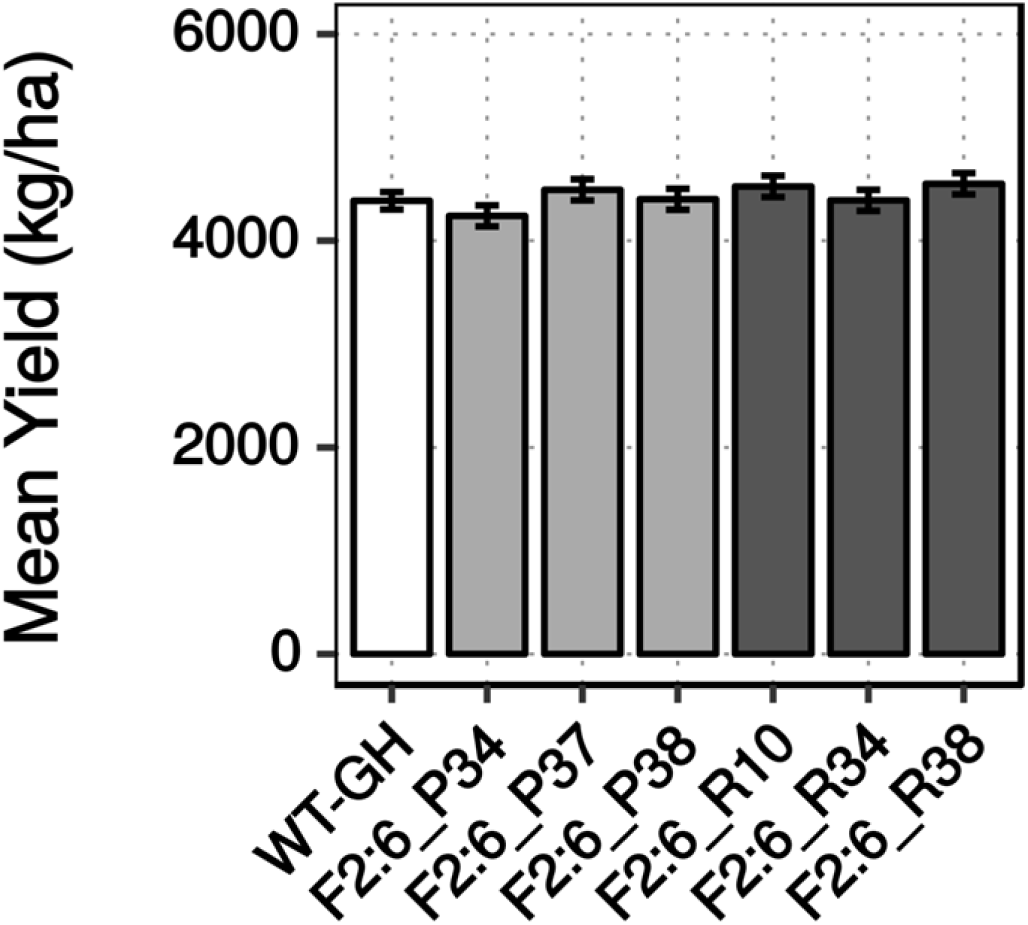
Bar graph showing reduction in *MSH1*-derived enhanced growth in epi F_2_:6. Bar graph showing yield measured as total seed weight in kg/ha normalized to 13% moisture from F_2_:6 lines compared to wild type (data pooled from three replicates each from four locations).

**Figure S5.**
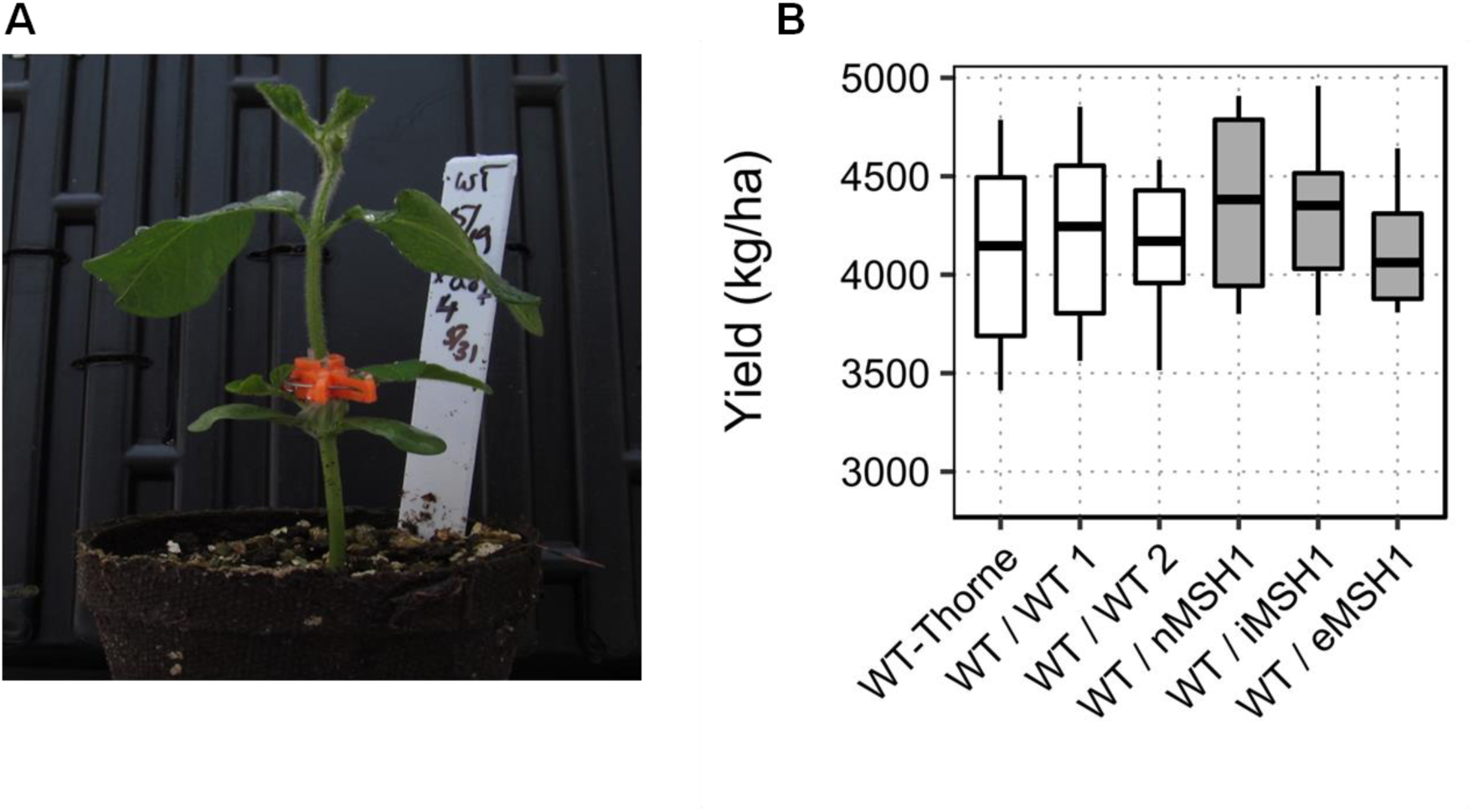
*MSH1*-derived enhanced growth in S2 progenies of wild type scion grafted onto *MSH1*-RNAi and *msh1* memory rootstock. **A)** Photo of a graft from a 10-day-old wild type scion grafted on 12-14 day-old *MSH1*-RNAi rootstock using wedge grafting method. **B)** Whisker plot of yield in S2 progenies from grafts measured as total seed weight in kg/ha corrected to 13% moisture (data pooled from three replicates each from four locations).

**Figure S6.**
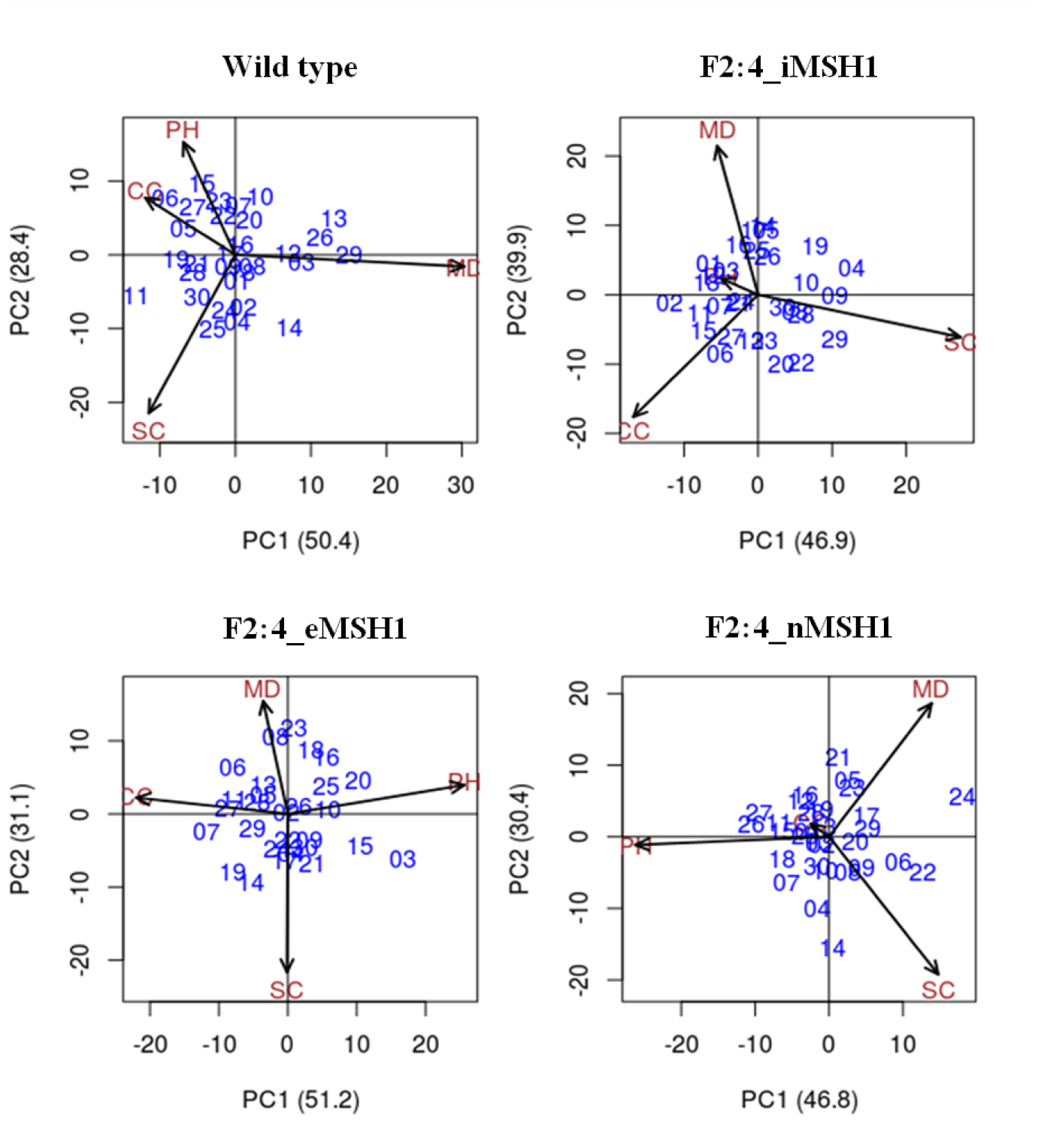
AMMI plots showing grouping of epi-F_2_:4 and wild type sub-lines across four different environments. Additive Main Effects Multiplicative Interactions (AMMI) plots using principal component analysis to partition the e x E interaction in wild type and epi-populations.

**Figure S7.**
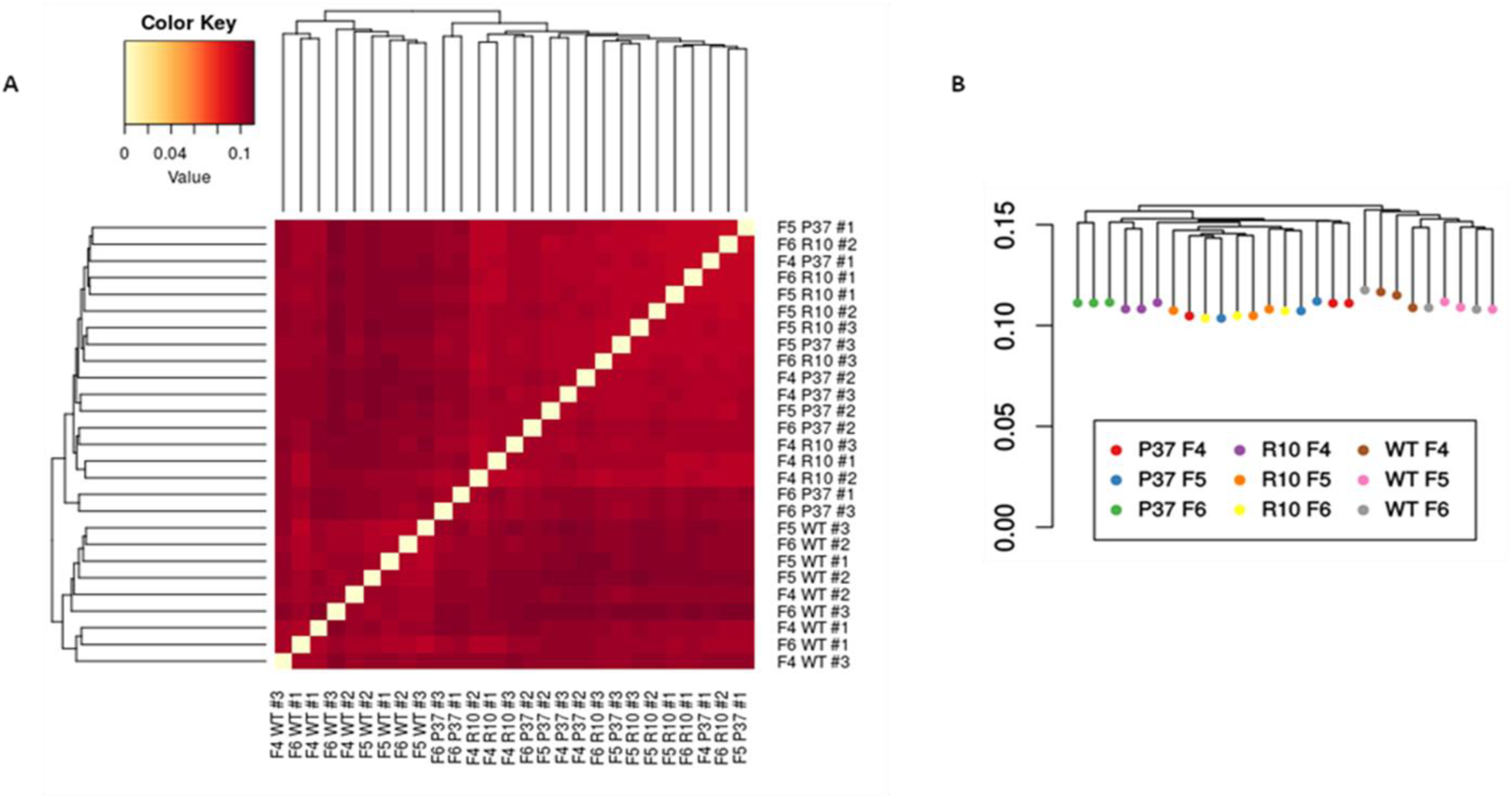
Genetic distance profiles using SNPs from transcriptome data of wild type and epi-lines. **A)** SNP profiles created from RNAseq dataset using the GATK toolkit for wild type and epi-lines showing no evidence of genetic contamination / unintentional hybridization in the enhanced growth epi-lines. **B)** Genetic dissimilarity plot confirming lack of evidence for genetic contamination in the experimental lines

**Figure S8.**
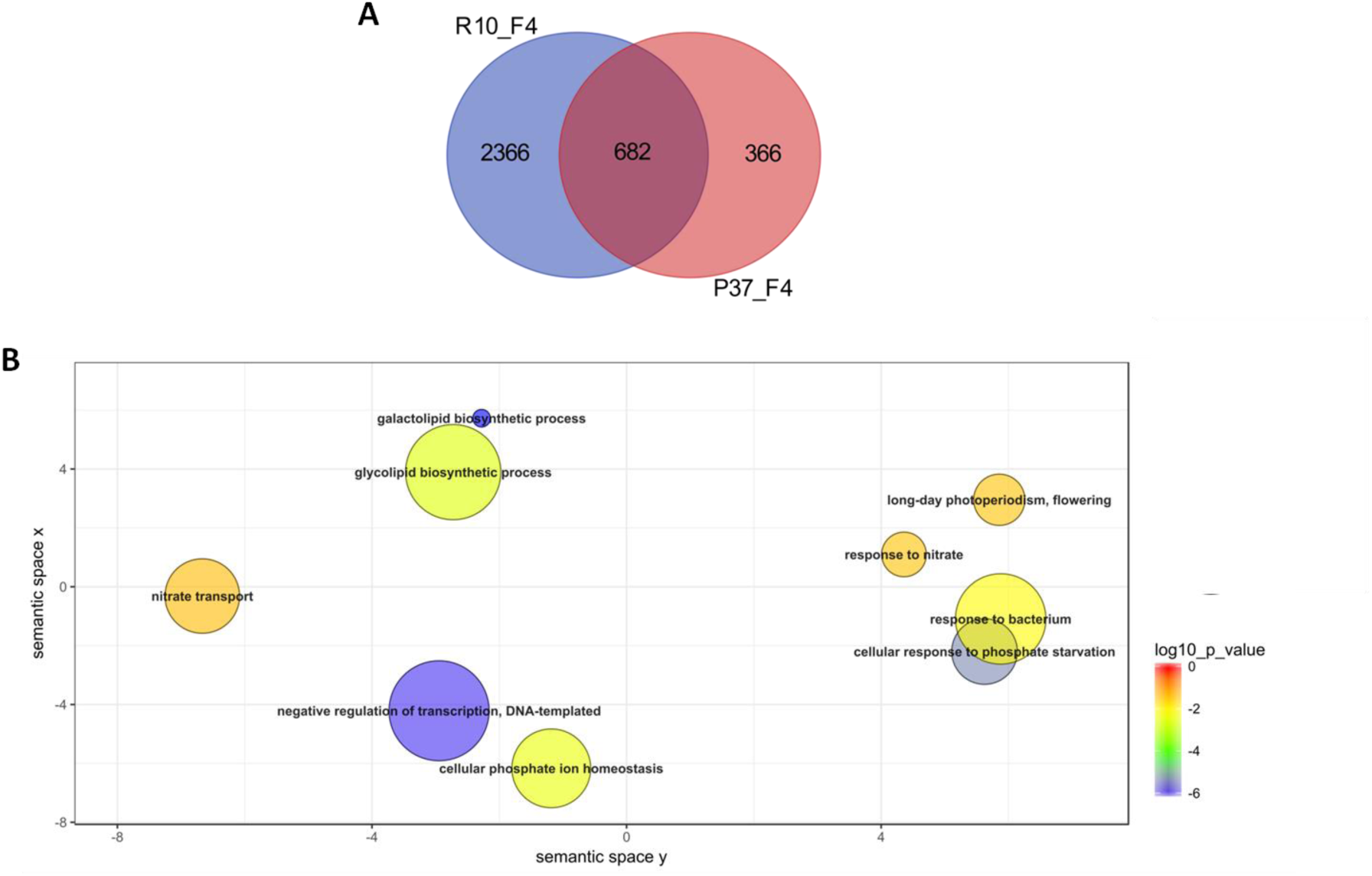
Overlap of genes and associated pathways in two epi-lines R-10 and P-37 with enhanced growth. **A)** Venn diagram showing overlapping DEGs between enhanced growth epi-line R-10 F_2_:4 and P-37 F_2_:4 (p-value < 0.05 and |log2(value)| > 0.5). **B)** REVIGO terms associated with 682 genes common between R-10 F_2_:4 and P-37 F_2_:4. GO terms (p-value < 0.05) obtained from SoyBase were used for REVIGO analysis using default parameters in agriGO.

